# Intermolecular interactions drive protein adaptive and co-adaptive evolution at both species and population levels

**DOI:** 10.1101/2021.02.08.430345

**Authors:** Junhui Peng, Nicolas Svetec, Li Zhao

**Affiliations:** Laboratory of Evolutionary Genetics and Genomics, The Rockefeller University, New York, NY 10065, USA

## Abstract

Proteins are the building blocks for almost all the functions in cells. Understanding the molecular evolution of proteins and the forces that shape protein evolution is essential in understanding the basis of function and evolution. Previous studies have shown that adaptation frequently occurs at the protein surface, such as in genes involved in host-pathogen interactions. However, it remains unclear whether adaptive sites are distributed randomly or at regions associated with particular structural or functional characteristics across the genome, since many proteins lack structural or functional annotations. Here, we seek to tackle this question by combining large-scale bioinformatic prediction, structural analysis, phylogenetic inference, and population genomic analysis of *Drosophila* protein-coding genes. We found that protein sequence adaptation is more relevant to function-related rather than structure-related properties. Interestingly, intermolecular interactions contribute significantly to protein adaptation. We further showed that intermolecular interactions, such as physical interactions may play a role in the co-adaptation of fast-adaptive proteins. We found that strongly differentiated amino acids across geographic regions in protein-coding genes are mostly adaptive, which may contribute to the long-term adaptive evolution. This strongly indicates that a number of adaptive sites tend to be repeatedly mutated and selected in evolution, in the past, present, and maybe future. Our results highlight the important roles of intermolecular interactions and co-adaptation in the adaptive evolution of proteins both at the species and population levels.

## Introduction

Natural selection plays an important role in the molecular evolution of protein sequences. Recent advances in genome sequencing and reliable inference methods at both phylogenetic and population levels have enabled fast and robust estimation of evolutionary rates and adaptation driven by natural selection. In addition, the increased availabilities of structural and functional data of proteins have made it possible to study how structural and functional constraints affect protein sequence evolution and adaptation. Different proteins and different sites within a protein have varying rates of evolution and adaptation due to both structural and functional constraints (Echave et al., 2016; Kosiol et al., 2008; Lindblad-Toh et al., 2011; Zhang and Yang, 2015). For example, genes that are highly expressed or perform essential functions are often under strong purifying selection and tend to evolve slowly (Drummond et al., 2005; Moutinho et al., 2019; Pál et al., 2001; Zhang and He, 2005; Zhang and Yang, 2015); genes involved in host-pathogen interactions, e.g., immune responses and antivirus responses, show exceptionally high rates of adaptive changes (Enard et al., 2016; Nielsen et al., 2005; Obbard et al., 2009; Palmer et al., 2018; Sackton et al., 2007; Sironi et al., 2015; Uricchio et al., 2019); and residues that are intrinsically disordered or at the protein surface are fast evolving and proved to be hotspots of adaptive evolution (Afanasyeva et al., 2018; Goldman et al., 1998; Lin et al., 2007; Moutinho et al., 2019; Ramsey et al., 2011). More recently, Slodkowicz & Goldman (Slodkowicz and Goldman, 2020) employed genomic-scale integrated structural and evolutionary phylogenetic analysis in mammals and showed that positively selected residues are clustered near ligand binding sites, especially in proteins that are associated with immune responses and xenobiotic metabolism. However, it remains unclear how adaptive sites are distributed in the genome and how adaptation is related with functions and structures. Moreover, most of the existing literature is focused on protein differences between species, and it remains unclear how much of within-species selective processes like spatially varying selection may contribute to long-term evolution.

Although evidence has shown that adaptation is more likely to occur at intrinsically disordered regions (Afanasyeva et al., 2018) and clustered at the surface of proteins (Dasmeh et al., 2013; Moutinho et al., 2019; Slodkowicz and Goldman, 2020), it remains unclear how functional and structural properties of proteins shape adaptation at the species and population scale. Moreover, due to the lack of structural and functional information of many proteins in the genome, the underlying evolutionary mechanism derived from current studies might be incomplete. Here, we systematically investigated the evolution and adaptation of protein-coding genes in *Drosophila melanogaster* by comparing it to its closely related species and their own populations, to distinguish the main factors that impact the evolution and adaption at the protein-coding level. We applied large-scale bioinformatic and structural analysis to obtain the structural and functional properties of proteins. We then classified residues into different structural and functional sites. By comparing rates of sequence evolution and adaptation between different proteins and sites, we were able to locate hotspots of adaptation at the genome scale. We found that functional properties are better predictors of protein adaptation rates than structural properties. Interestingly, we found that adaptation rates of a protein positively correlate with the fraction of residues that are involved in intermolecular interactions inside the protein. In agreement with this finding, we found that putative binding regions including allosteric sites at protein surface show higher rates of adaptive evolution than other sites. For proteins under fast-adaptive evolution, we showed that they tend to interact with each other more frequently than random expectations, suggesting fast-adaptive genes might undergo co-adaptive evolution. We further discovered that co-adaptation might be universal for many interacting proteins in *D. melanogaster*.

Our results suggest that intermolecular interactions in *D. melanogaster* are an important driver of protein adaptive evolution. We further hypothesized and provided evidence that intermolecular interactions, such as physical interactions might be an important mechanism that contribute to the co-adaptive evolution of interacting proteins in *D. melanogaster* genome. One of the intriguing questions is that how the adaptive signals in populations (short-term) and between species (long-term) are correlated. We then asked if those patterns hold for selective process occurring within species. Despite the abundance of literature studying geographic variation in Drosophila species (Bergland et al., 2014; Fabian et al., 2012; Kolaczkowski et al., 2011; Lack et al., 2015; Langley et al., 2012; Pitchers et al., 2013; Reinhardt et al., 2014; Svetec et al., 2016), very little is known for a systematic evaluation of the protein properties affected by spatially varying selection. We thus investigated protein adaptation signals of strongly differentiated amino acids across geographic regions, which were often associated with within-species local adaptations (Matthey-Doret and Whitlock, 2019). We showed that most of the patterns found in the between-species in fact hold at the within species levels. This may partly because most sites contributing to within-species local adaptation tend to also contribute to long-term adaptive evolution in *D. melanogaster*, suggesting that a subset of protein-coding loci are constantly or repeatedly utilized for adaptive purpose.

## Results

### Putative molecular interaction sites are hotspots for protein adaptive evolution

To uncover the main factors that impact the evolutionary rates of genes, we analyzed 13,528 protein-coding genes in *D. melanogaster* using genome data from *melanogaster* subgroup species and *D. melanogaster* population genomics data from 205 inbred lines from Drosophila Genetic Reference Panel, Freeze 2.0, DGRP2 (Huang et al., 2014). We applied a maximum likelihood method (Yang, 2007) to compute the dN/dS ratio (ω) using the protein-coding sequences of five closely related melanogaster subgroup species (*D. melanogaster*, *D. simulans*, *D. sechellia*, *D. yakuba* and *D. erecta*). We estimated the proportions of adaptive changes (α) in each gene by applying an extension of McDonald–Kreitman (MK) test named asymptotic MK (Messer and Petrov, 2013; Uricchio et al., 2019) using *D. yakuba* as outgroup. We then calculated the rate of adaptive changes (ω_a_) of each gene by multiplying ω to α (ω_a_ = αω) (Moutinho et al., 2019) (See *Material and Methods*). The rate of nonadaptive changes can be further calculated by ω_na_=ω-ω_a_. Finally, we successfully assigned ω to 12,118 protein coding genes and ω_a_ and ω_na_ to 7,192 genes. For each of *D. melanogaster* genes subjecting the same analysis pipeline, we further obtained 17 different structural or functional properties (see *Material and Methods*, supplementary file S1, Table S1-S2). We calculated Pearson’s correlations of ω, ω_a_ and ω_na_ with all these properties (Table S1). Many of these genome-wide correlations were expected (for details, see Supplement Information, Table S1-S2, Fig. S1-S5). Interestingly, we found that some previously unexplored properties, fractions of molecular-interaction sites (including PPI-site ratio, ratio of residues involved in protein-protein interactions, and DNA-site ratio, ratio of residues involved in protein-DNA interactions) were strongly positively correlated with ω, ω_a_, and ω_na_ (Supplement Information, section *Molecular interactions contribute to the variations of protein sequence evolution and adaptation*, Table S1, Fig. S1-S2). The results indicate that molecular interactions might act as an important factor that drives protein adaptive evolution in the *Drosophila* genome.

We then investigated whether residues involved in molecular interactions are targets for adaptive evolution. To tackle this question, we predicted protein-protein interaction sites (PPI-sites) and DNA binding sites (DNA-sites) for each of *D. melanogaster* protein sequences (see Material and Methods). In addition, we characterized allosteric residues as surface and interior critical residues with STRESS model (Clarke et al., 2016) for all the structural models. We also extracted putative binding sites from STRESS Monte Carlo (MC) simulations. We calculated ω, ω_a_, and ω_na_ for residues in each of the putative molecular interaction categories. Strikingly, we observed that residues involved in protein-protein interactions, DNA binding and ligand binding exhibited higher rates of adaptive evolution compared to their corresponding null sites (t-test, p = 7e-10, 0.18 and 7e-15, respectively) (Fig. 1A-C). In addition, allosteric residues at protein surface showed higher adaptation rates than allosteric residues at protein interior (t-test, p = 3e-10) or residues that are not involved in ligand binding (t-test, p = 0.003) (Fig. 1C, Fig. S6).

**Fig. 1.**
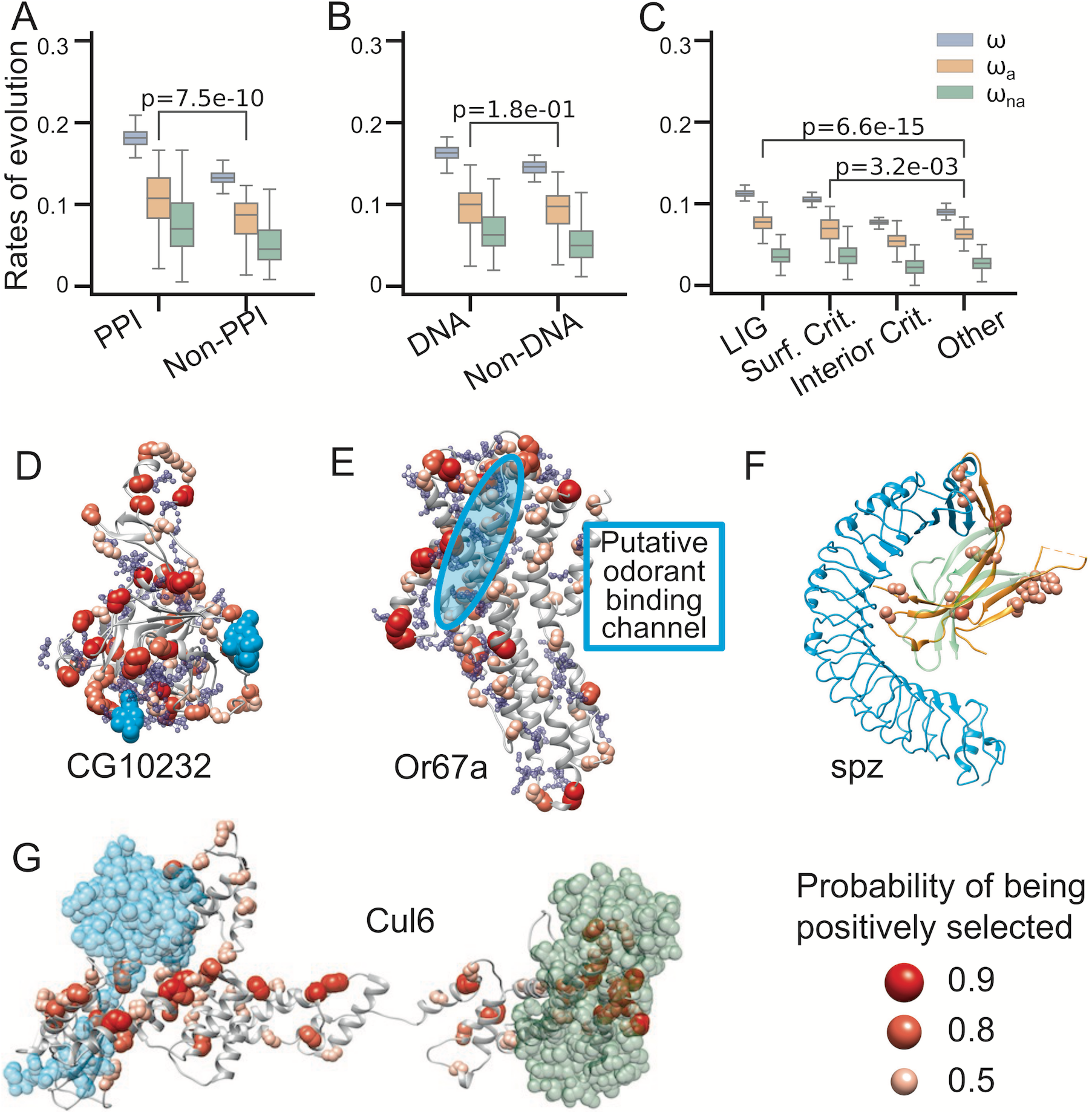
Adaptive evolution in molecular interaction sites. Protein-protein interaction sites (A), DNA binding sites (B) and putative ligand binding sites (C) show higher adaptation rates than none binding sites. The t-test p-values between the adaptation rates of binding sites and non-binding sites were highlighted in (A), (B) and (C). T-test p-values between other evolutionary rates of binding sites and non-binding sites were shown in Fig. S2. Examples of positive selection around molecular interaction sites in high quality structural models of CG10232 (D), Or67a (E), spz (F), and Cul6 (G). Except for spz (PDB code 3e07), the other proteins are obtained from SWISS model repository. Putative ligand binding pockets of CG10232 (D) and Or67a (E) are shown in blue spheres. Ligands including interacting proteins are shown in cyan or green: NAG of CG10232 in cyan (D), Toll receptor of spz in cyan (F), RING-box protein in cyan and F-box protein in green for Cul6 (G). The putative odorant binding channel of Or67a is highlighted in cyan circle (E). The ligand poses in (D, F and G) are obtained by superimposition from structure 2XXL, 4BV4 and 1LDK, respectively.

To gain a better understanding of adaptation in molecular interaction sites, we further visualized positive selections that are associated with molecular interactions. We first investigated whether adaptive evolution is associated with particular protein structures or protein families. To do this, we looked into fast-adaptive proteins with the largest ~15% rates of adaptation (ω_a_ > 0.15) that are linked to high-quality structural models. Interestingly, among these proteins, we found 45 enriched as trypsin-like cysteine/serine peptidase domain and 17 7TM chemoreceptors, suggesting widespread adaptive evolution acting on these protein families or protein domains in *D. melanogaster* (Table S3). Many of the 7TM chemoreceptors are olfactory and gustatory genes and show adaptive evolution in various species such as *Drosophila* and mosquito (Hill et al., 2002; Lawniczak and Begun, 2007; McBride, 2007; Wu et al., 2009). In addition to these two protein families, previous studies identified recurrent positive selections acting on some other fast-adaptive proteins in *Drosophila* and mammals, and the possible adaptive evolution mechanisms have been linked to exogenous ligand binding, for example, serine protease inhibitors (serpin), Toll-like receptor 4 (TLR-4), and cytochrome P450 (Jiggins and Kim, 2007; Slodkowicz and Goldman, 2020).

We used the two representative cases of fast adaptive protein evolution of CG10232 and Or67a - a trypsin-like cysteine/serine peptidase domain and a 7TM chemoreceptor, respectively - to illustrate the link between adaptive evolution and molecular interactions in the two protein families with frequent adaptive evolution. We observed that in both cases, positively selected sites were significantly closer to predicted or inferred binding sites in the protein 3D structure (t-test, p = 8e-133 for CG10232 and 3e-169 for Or67a, Fig. 1D-E) and were overlapped with predicted or inferred binding pockets for CG10232 (Fisher’s exact test, p-values 0.02, Fig. 1D). There might be an overlap with predicted or inferred binding pockets for Or67a, but it is not statistically significant (Fisher’s exact test, p = 0.19, Fig. 1E). Specifically, in CG10232, we found clusters of positive selected sites around NAG binding sites that are inferred from a crystal structure of serine protease (PDB code: 2XXL) (Fig. 1D), while in Or67a, positively selected sites expand around the putative odorant binding channel formed by helices S1-S6 in extracellular regions (Butterwick et al., 2018) (Fig. 1E).

Besides the examples that are associated with exogenous ligand or exogenous peptide binding, we also identified two previously undescribed examples where adaptive evolution might be linked to endogenous protein binding: Spaztle (spz, Fig. 1F) and Cul6 (Fig. 1G). Spaztle can bind to Toll-like receptors (TLR) and trigger a humoral innate immune response. We built the missing loop in Spaztle in the crystal structure of Toll/Spaztle complex (PDB code 4BV4) according to the dimeric crystal structure of Spaztle (PDB code 3E07). In this complex structural model, we observed several positively selected sites in Toll-4/Spaztle interfaces (Fig. 1F). Cul6, another example, is a protein in the cullins family in *D. melanogaster*. The cullins protein family are known as scaffold proteins that assemble multi-subunit Cullin-RING E3 ubiquitin ligase by forming SCF complex with F box and RING-box (Rbx) proteins (Zheng et al., 2002). We constructed the putative Cul6 contained SCF complex by superimposition to the crystal structure of the Cul1-Rbx1-Skp1-F box^Skp2^ SCF ubiquitin ligase complex (Zheng et al., 2002). In the structural model, we observed positively selected sites in Cul6 clustered around the binding sites of RING-box protein, Rbx1, and F-box protein, Skp1 (Fig. 1G). The examples above suggest that intermolecular interactions, including both exogenous and endogenous binding, could contribute to protein adaptive evolution.

### Frequent adaptive evolution and co-adaptative evolution in genes involved in reproduction, immune system, and environmental information processing

To find out whether specific biological functions were associated with fast-adaptive genes, we applied DAVID GO analysis to the genes with the largest rates of adaptation (ω_a_ > 0.15, top ~15%). The significant GO terms are frequently linked to serine-type endopeptidase activity, reproduction, protein lysis, chemosensory and other related biological functions (Table S4). As these fast-adaptive genes tend to be enriched in similar biological functions, we asked whether these genes evolved co-adaptively, i.e., whether these proteins are interacting with each other frequently. To test this possibility, we obtained PPI of *D. melanogaster* from STRING database (Szklarczyk et al., 2019) and analyzed protein-protein interactions among fast-adaptive proteins. We found that fast-adaptive proteins tend to interact with each other more frequently than expected (PPI enrichment p-value < 1.0e-16). In the PPI network of fast-adaptive proteins, we observed 7 strongly connected sub-clusters with at least 5 members (Fig. 2A, Table S5, example Fig. 2B-C). Proteins in these sub-clusters are enriched in biological processes such as reproduction, immune response, defense response to bacterium and virus, RNA interference, chitin metabolic, etc., (Table S5-S11) which are in line with the GO analysis of fast-adaptive genes (Table S4), and previous enrichment analysis of positively selected genes identified from genome-wide studies (Begun et al., 2007; Enard et al., 2016; Nielsen et al., 2005).

**Fig. 2.**
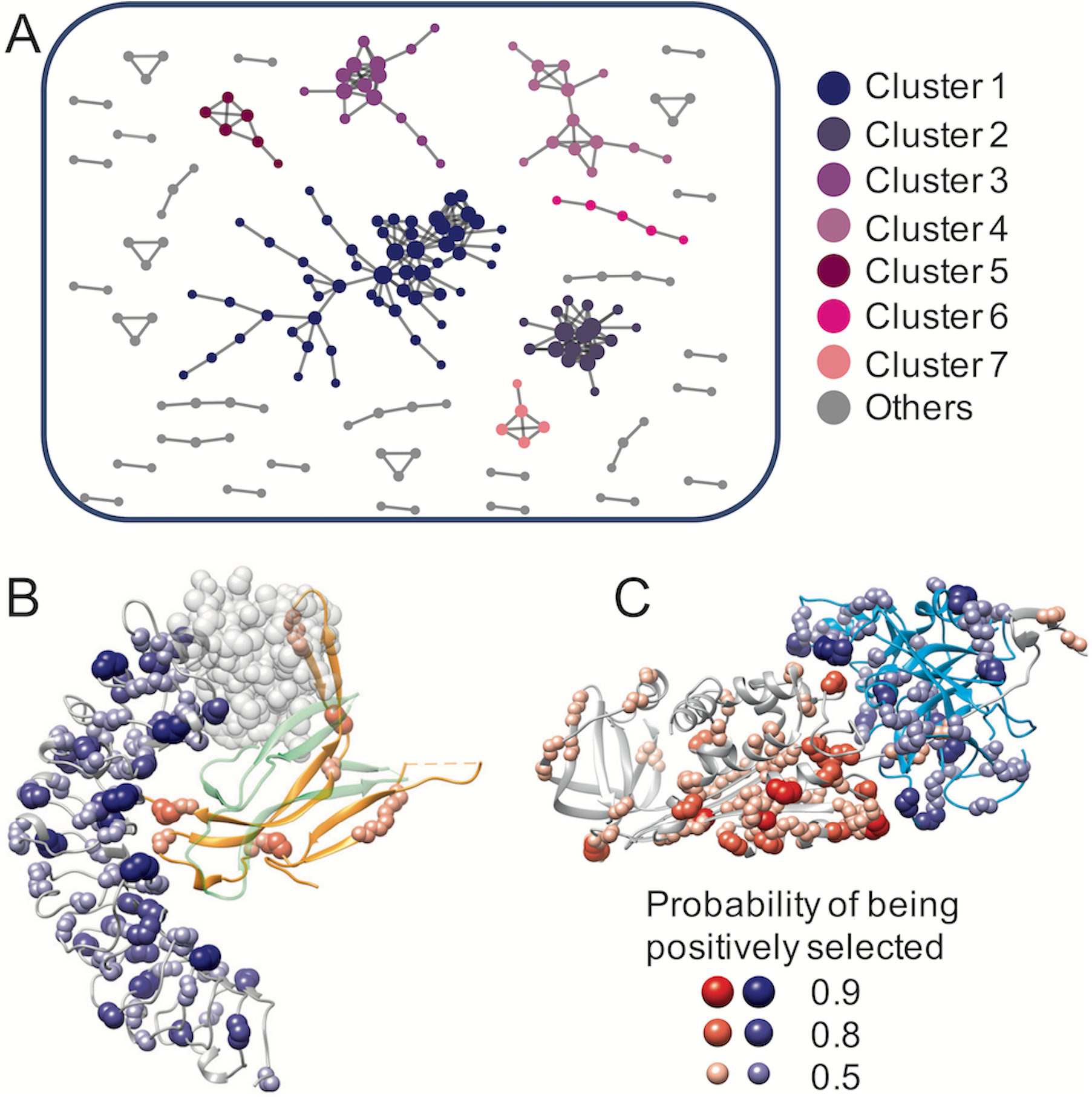
Co-adaptation of fast-adaptive proteins. (A) Sub-clusters of PPI networks of fast-adaptive proteins. Only proteins with at least one partner were shown. Examples of molecular interactions that might regulate co-adaptation in fast-adaptive proteins: (B) Toll-4 (gray) and spz (orange, with green representing the other spz monomer), (C) Spn28Db (gray, serine protease inhibitor 28Db) and CG18563 (cyan, with Go term “serine-type endopeptidase activity”). A putative N-terminus (transparent beads) of Toll-4 were built by superimposition from 4LXR, since the N-terminus were missing in the structural model. Complex structural model of Spn28Db and CG18563 was inferred from 1EZX.

We next asked whether co-adaptation plays a role in the adaptive evolution of interacting proteins to a broader extend, including both fast- and slow-adaptive proteins. To address this question, we analyzed and compared adaptation rates of all D. *melanogaster* PPIs available in STRING database with high confidence, and we found that protein partners of fast-adaptive proteins (ω_a_>0.15) have significantly larger maximum/average ω_a_ compared to slow-adaptive proteins (Fig. 3). We further analyzed and visualized adaptive evolutionary rates of proteins in PPI networks of 9 different biological pathways extracted from KEGG pathways, including the immune system, xenobiotics biodegradation, response to the environment, aging and development, genetic information processing, sensory system, transport and catabolism, cell growth and death and metabolism. We observed that, in these PPI networks, proteins with relatively large ω_a_ tend to interact with each other (Fig. 4A-B). We also noticed that, for pathways that are previously known as adaptation-hotspots (e.g., immune system), fast-adaptive proteins can act as central nodes and are co-adaptively evolving with other fast-adaptive proteins (Fig. 4A, C). While in pathways such as transport and catabolism, fast-adaptive proteins are mainly at PPI periphery. Specifically, we observed that in pathways related to immune system and environment adaptation, where adaptive evolution often occurs, fast-adaptive proteins have comparable number of interactors as other genes (Fig. 4C-D), while in conserved pathways such as transport and catabolism, fast-adaptive genes are often at network peripheries and have significantly fewer interactors (Fig. 4E-F). In line with these findings, we found that ω_a_ are larger in pathways that harbor fast-adaptive proteins as central nodes than other pathways (Fig. S7).

**Fig. 3.**
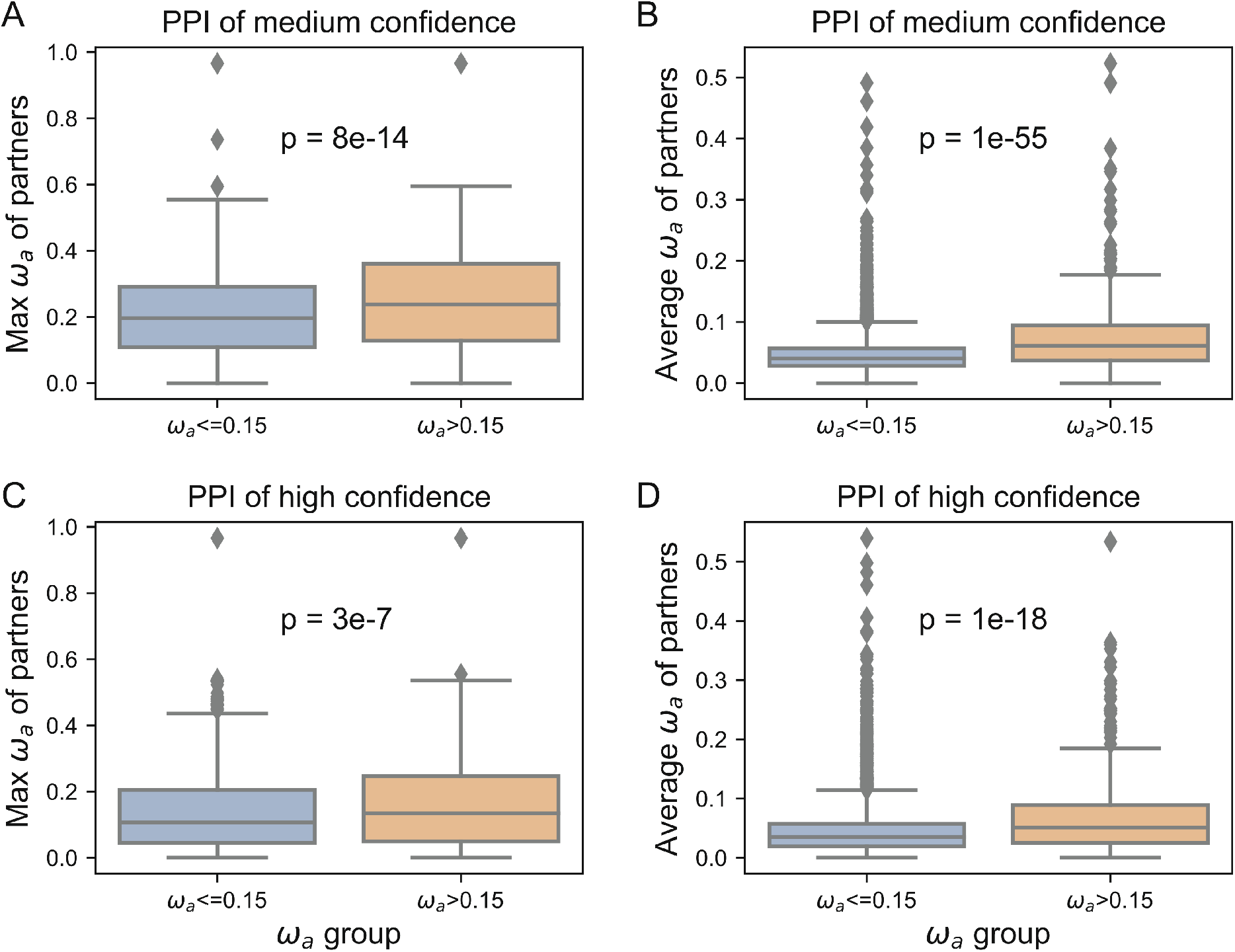
Co-adaptation of PPIs in *D. melanogaster*. For fast-adaptive proteins, adaptation rates of their partners (orange box plot) are significantly larger compared to slow adaptive proteins (blue box plot). Max ω_a_ of protein partners are shown in (A and C) and averaged ω_a_, of protein partners are shown in (B and D). PPI from STRING with median confidence (combined score larger than 0.4) are shown in (A and B), and PPI with high confidence (combined score larger than 0.7) are shown in (C and D).

**Fig. 4.**
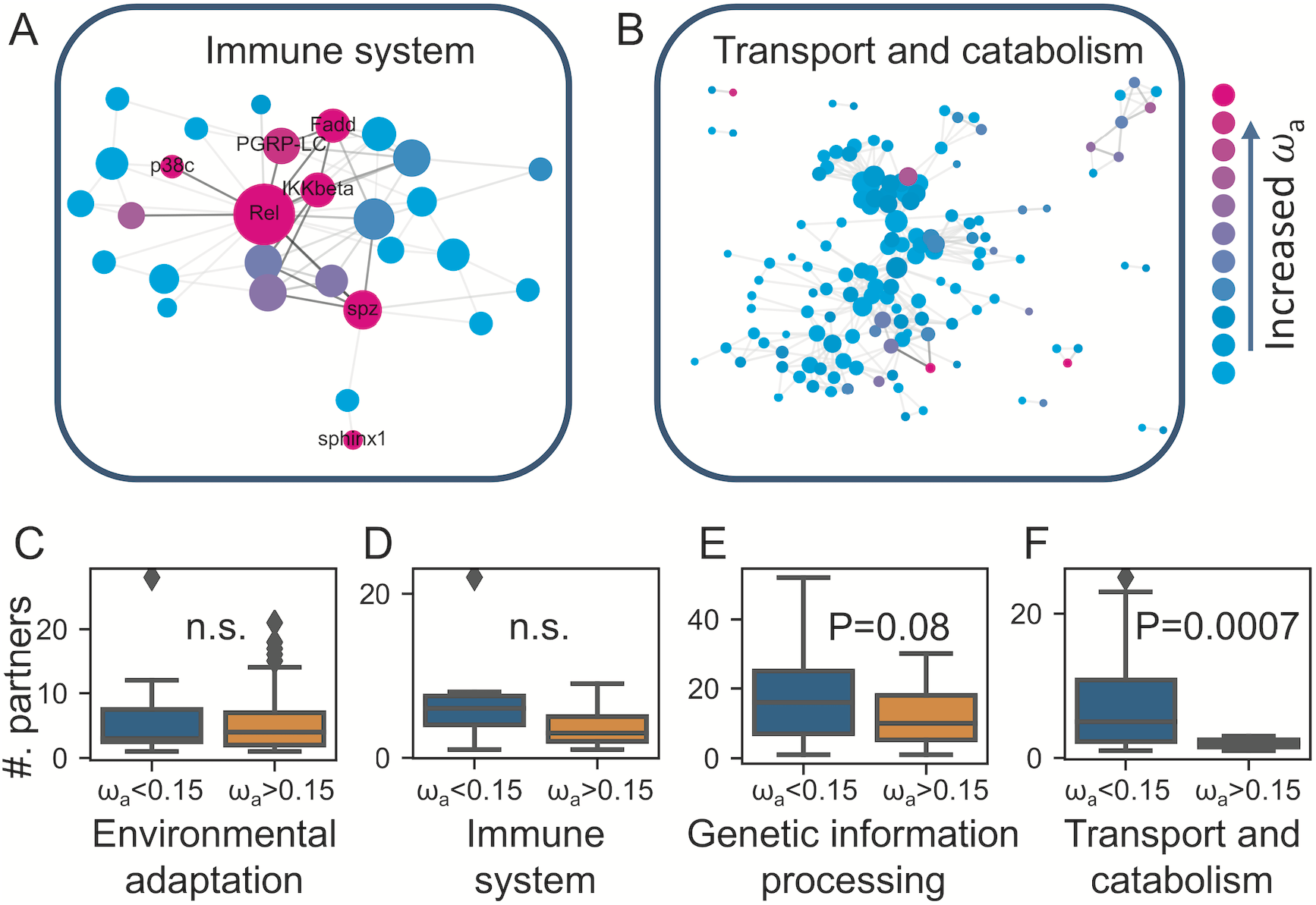
Rates of protein sequence adaptive evolution in the PPI network of different functional pathways. The PPI networks showed the adaptive evolution in immune system (A) and transport and catabolism (B). In pathways that are hotspots of adaptive evolution, e.g., environmental adaptation (C) and immune system (D), fast-adaptive proteins can act as central nodes. While in conserved pathways, e.g., genetic information processing (E) and transport and catabolism (F), fast-adaptive proteins are often at the periphery of the PPI network.

### Physical interactions contribute to co-adaptation of fast-adaptive genes

Having established that molecular interactions contribute to the adaptive evolution of protein sequence, we then investigated whether these physical molecular interactions could drive protein-protein co-adaptation. To do this, we looked into interacting fast-adaptive protein pairs that are associated with known or inferred complex structural models. For inferred complex structural models, we superimposed the structural models of the pair of proteins onto their high resolution homologous complex structures. Here we illustrated co-adaptation at PPI interface in two examples: Toll-4/Spatzle and Spn28Db/CG18563 (Fig. 2B-C).

#### Toll-4/Spatzle

Toll-4 is a member of toll-like receptors. Previous studies have shown strong evidence of adaptive evolution of Toll-4 in *Drosophila* and mammals (Levin and Malik, 2017; Slodkowicz and Goldman, 2020), and implied the important roles of ligand binding in the positive selection and coevolution of Toll receptors and Spatzle proteins (Lima et al., 2021). Toll-4 can bind to Spatzle and trigger further innate immune responses with high confidence (inferred from STRING database). In the previous section, we showed that several positively selected sites in Spatzle overlap with Toll-Spatzle interfaces (Fig. 1F). Here, we further showed that, in Toll-4, many significantly positively selected sites were located at interface for Spatzle (Fig. 2B), which is in line with a previous study of Toll-4 in *D. willistoni* (Levin and Malik, 2017).

#### Spn28Db/CG18563

Spn28Db is one of the serine protease inhibitors in *D. melanogaster* expressed in male accessory glands, while CG18563 belongs to the protein family of trypsin-like cysteine/serine peptidase domain. The interactions between the two proteins were predicted with high confidence from the STRING database, and the molecular interactions can be inferred from existing crystal structure of serpin and bacteria protease complex (PDB code 1EZX). We observed many positive selected sites at the molecular interface between the two proteins (Fig. 2C), suggesting that physical interactions might play a role in the co-adaptation of the two proteins.

### Most geographically differentiated non-synonymous SNPs in protein-coding genes are adaptive

To learn more about the relationship between short-term adaptation to local environments and long-term adaptive evolution, we extracted residues with significant allele frequency differentiation across latitudes in North America (Svetec et al., 2016) and Africa (Lack et al., 2015). For the DPGP3 African population data, we followed the same protocol as Svetec et al. 2016 to identify significantly differentiated SNPs (section *Population genetics of DPGP3 African population* in Material and Methods). We then computed evolutionary rates (ω), adaptation rates (ω_a_), non-adaptation rates (ω_na_) and proportions of adaptive changes (α) of these residues as in the previous section (Fig. 5A-B, Fig. S8). We observed that, in both the North American population and the African population, these sites have significantly higher proportions of adaptive changes (Fig. 5A-B) than other SNPs, suggesting that they can be hotspots for adaptive evolution. To find out whether these SNPs are related to even longer-term adaptive evolution, we inferred positive selection sites of each protein-coding gene from phylogenic data (see Material and Methods). We found that geographically differentiated non-synonymous SNPs are significantly enriched for long-term positive selection (Fig. S9). To further characterize structural and functional properties of short-term genetic variations, we mapped geographically differentiated non-synonymous residues to different structural and functional characteristics, such as ISD, RSA, PPI-sites, DNA-sites and ligand-binding sites. We found that these non-synonymous SNPs were significantly enriched in disordered regions and protein surfaces, as well as in protein-protein interactions and ligand-binding (Fig. S9). To better visualize the characteristics of these SNPs, we used *Toll-4* as an example. We mapped its geographically differentiated non-synonymous sites onto its structural model. We found that these sites are either positively selected or are located very close to positively selected sites (Fig. 5C-D). For example, highly differentiated sites in North American population, N279 (FDR 3e-7) and H431 (FDR 3e-6) were predicted to be positively selected both at a probability of p=0.9. While another highly differentiated site, D424 was close to three positively selected sites S401 (p=0.8), H431 (p=0.95) and V448 (p=0.8). We also noticed some differentiated sites that may be located within ligand binding sites, including F297 (FDR 3e-3), S311 (FDR 3e-3), H431 (FDR 3e-6) and H462 (FDR 1e-2). In the structure of Toll4, we also observed 3 highly differentiated sites in African populations, which are S311 (FDR 4e-2), H431 (FDR 2e-2) and S490 (FDR 2e-4). We noticed that all the 3 sites overlapped with differentiated SNPs in North America and two of them (S311 and H431) localized within ligand binding sites (Fig. 5D), further supporting our observation that ligand binding sites in some genes may undergo recurrent adaptive evolution.

**Fig. 5.**
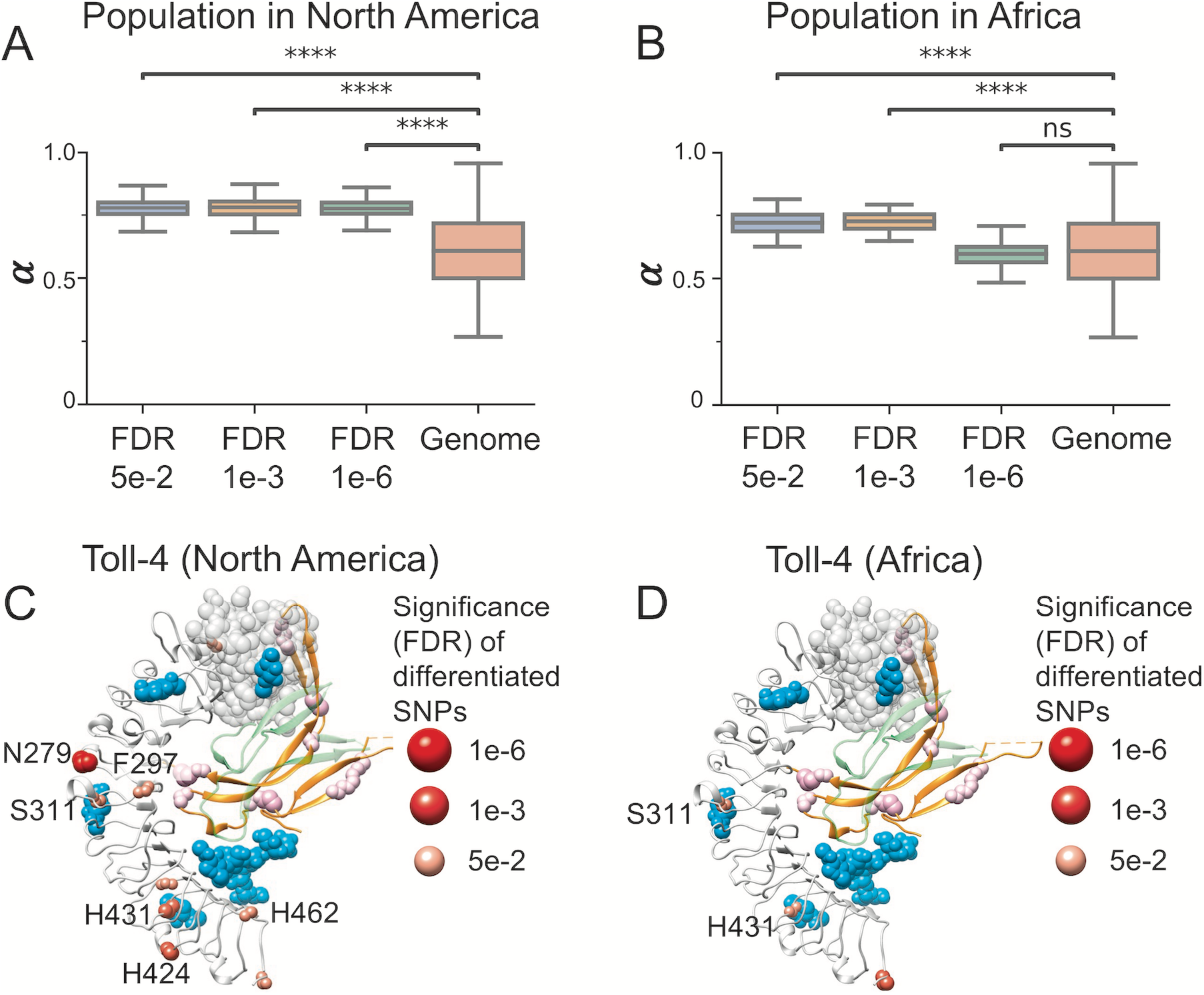
Adaptive evolution in significantly differentiated SNPs. The significantly differentiated SNPs at different FDR cutoffs all show much higher proportions of adaptative changes (α) than genome-wide expectation in North American population (A) and African population (B) (t-test_ind, ****, p<1e-4). (C) Significant differentiated nonsynonymous SNPs of North American populations in Toll-4. Ligands are shown in cyan by superimposing crystal structure of Toll-Spatzle (PDB code 4BV4) on to Toll-4 structural model. Residues N279 and H431 are both highly differentiated (FDR 3e-7 and 3e-6) and positively selected (both at probability of p=0.9). Other highly differentiated sites, F297, S311, H424, H431 and H462 are located near ligand binding sites or positively selected sites. (D) Significantly differentiated nonsynonymous SNPs of African populations in Toll-4. Two highly differentiated SNPs, S311 (FDR 4e-2) and H431 (FDR 2e-2) exist in North American population and are located near ligand binding sites.

## Discussion

In this study, we systematically studied the impact of structure- and function-related gene properties on protein sequence evolution and adaptation in *D. melanogaster* genome. We found that molecular interactions in proteins contribute to the variation of protein sequence adaptive evolution. A novel discovery of this work is that molecular interaction sites including protein-protein interaction sites and protein-DNA interaction sites are hotspots for adaptative evolution. We revealed that fast-adaptive proteins tend to interact with each other frequently and protein partners of these fast-adaptive proteins tend to have higher adaptation rates, suggesting that co-adaptive evolution might be common in *D. melanogaster*. By visualizing examples of interacting fast-adaptive proteins, we further demonstrated that physical interactions may contribute to the co-adaptation of fast-adaptive proteins.

Protein surface and intrinsic disorder regions are frequent targets for adaptive evolution and contribute to the variations of protein sequence adaptive evolution (Afanasyeva et al., 2018; Moutinho et al., 2019). However, the detailed mechanisms underlying these observations remains unclear. One possible explanation would be that these regions are frequently linked to intermolecular interactions (Afanasyeva et al., 2018; Moutinho et al., 2019). For example, Moutinho et al. hypothesized that molecular interactions involved in host-pathogen coevolution were the major driver of protein adaptation (Moutinho et al., 2019). Here, we further identified that proportions of possible molecular interaction sites inside proteins contribute to the variations of protein sequence adaptive evolution. These molecular interaction sites or regulatory sites at protein surface can be hotspots of protein adaptation. Indeed, some specific molecular interactions have been linked to adaptive evolution in several case studies (Bachtrog, 2008; Hughes and Nei, 1988; Levin and Malik, 2017; Schott et al., 2014), a recent study on sweet taste receptors in the songbird radiation (Toda et al., 2021), and large-scale studies based on proteins with high quality structural models (Slodkowicz and Goldman, 2020). In the latter study, the authors showed that amino acids under positive selection in mammals tend to cluster closer to binding sites of exogenous ligands than expected by chance (Slodkowicz and Goldman, 2020), suggesting an important role of functionally important regions in adaptive evolution. Here, we extend the conclusion *to D. melanogaster* genome, including proteins with or without high resolution structural models. We also showed that in addition to exogenous ligands, endogenous ligands might also contribute to adaptive evolution, while the latter might explain why interacting proteins tend to evolve co-adaptively.

Notably, previous studies showed that multi-interface proteins tend to be evolving more slowly than single-interface proteins (Kim et al., 2006) and that proteins with many interactors tend to evolve slowly (Jordan et al., 2003), which seems to be contradictory to our results that proteins with more interaction sites evolve faster and have faster adaptation rates. Here, we argue that, in our study, we used sequence profiles to predict molecular interaction sites in proteins at a genomic scale, rather than only looking into proteins with high resolution structures. In this way, we may capture many weak or transient interactions, which are evolving faster than obligate and conserved interactions (Mintseris and Weng, 2005). Meanwhile, we did not exclude intrinsically disordered regions (IDR) or intrinsically disordered proteins (IDP) in our study, which are widespread in *D. melanogaster* genome. It has been suggested that IDR/IDP tend to evolve fast due to the lack of structural restraints (Echave et al., 2016). In the functional aspect, IDR/IDP are thought to be promiscuous binders through many multiple binding mechanisms, including forming static, semi-static, and fuzzy or dynamic complexes (Uversky, 2019), suggesting that the evolution of IDR/IDP cannot be explained merely by the lack of structural restraints. Indeed, IDP and IDR in the human genome were found to be undergoing extensive adaptive evolution (Afanasyeva et al., 2018). At last, it has been recognized that, except for allosteric regulations, encounter complexes (Gabdoulline and Wade, 1999) might also play an important role in mediating intermolecular interactions, such as protein-protein association (Tang et al., 2006) and protein-ligand binding (Re et al., 2019). Since encounter residues that are responsible for encounter complexes do not reside in conserved binding interfaces, these residues could be under relaxed purifying selection or even positive selection, which could be another yet-to-identify mechanism that contributes to protein sequence adaptive evolution.

We showed that fast-adaptive proteins are enriched in molecular functions such as reproduction, immunity, and environmental information processing (Begun and Lindfors, 2005; Begun and Whitley, 2000; Lazzaro et al., 2004). We further demonstrated that fast-adaptive proteins tend to interact with each other more frequently than random expectations, suggesting co-adaptation might be common among fast-adaptive proteins. Mechanisms contributing to the co-adaptation could be: (1) interacting fast-adaptive proteins are often enriched in similar molecular functions and under similar selective pressure; (2) interacting fast-adaptive undergo co-evolution through physical interactions. In this study, we showed two examples that adaptive evolution could occur at protein-protein interface, which suggest that physical interactions could contribute to the co-adaptation of fast-adaptive proteins in *D. melanogaster*. Moreover, we showed that co-adaptation might exist to a broader extend rather than only among fast-adaptive proteins. Specifically, proteins that interact with fast-adaptive proteins tend to have higher adaptation rates. Since molecular interactions contribute to adaptive evolution, it is reasonable to hypothesize that co-adaptation at a broader extend could be regulated by these interactions. Actually, it has been suggested that interacting proteins tend to have similar evolutionary rates and the possible mechanism would be the co-evolution of physical interactions (Pazos and Valencia, 2008).

In this study, we found that amino acids showing strong geographically differences often overlap with sites that show adaptive signals between species. These loci follow similar patterns as adaptive changes, i.e. they are enriched in disordered regions, protein surfaces, and functionally important regions. These results suggest that population differentiation of protein-coding genes can be an important basis for long-term adaptive evolution. In other words, many SNPs are repeatedly selected for the adaptive processes in evolution. Importantly, our results indicate that most of the clinal amino-acid changes are adaptive, suggesting that non-selective forces play a less essential role in the SNPs that show strong geographic differences. Our results also support a large effect of spatially varying selection on protein sequence and structures (Storz and Kelly, 2008). Interestingly, our previous work showed that geographically differentiated SNPs often occur on the same orthologous genes between species but rarely same SNPs (Zhao et al., 2015), it would interesting to extend this work to *D. simulans* to study parallel protein evolution at the structural level.

It should be noted that studies at the genomic scale that aim to uncover the function- or structure-related constraints imposed on protein sequence evolution and adaptation share similar limitations that for most of the proteins or residues, structural or functional information would be incomplete or even missing. To overcome this, in this study, we used highly accurate neural-network based tools to predict molecular interactions, secondary structures, intrinsic structural disorder, relative solvent accessibility for each of the proteins. In this way we were able to identify key factors that impact protein sequence evolution and adaptation in a less accurate but rather systematic fashion. Notably, it has been reported that false positives are nonnegligible in methods to the estimation of adaptive evolution (Markova-Raina and Petrov, 2011), and other mechanisms including translational selection were acting on evolution of protein-coding genes (Larracuente et al., 2008), which together hinder our understanding toward protein evolution and protein adaptation. Another limitation of our study is that we did not include indels, especially non-frameshift indels, as it is very difficult to address the adaptive effects of indels. However, with method development and increased knowledge of protein structures, this would be an important question to investigate in the future. In the recently years, deep learning-based predictors and estimators have contributed to our knowledge towards protein structure (Jumper et al., 2021), protein function (Kulmanov and Hoehndorf, 2020), or even protein evolution (Schrider and Kern, 2018). We hope that with the availability of more and more curated structural, functional information and complex structural models of proteins in the near future, we will be able to uncover the precise role of molecular interactions in protein sequence adaptive evolution.

## Material and Methods

### *d_N_*/*d_S_* ratio (ω)

We used a maximum likelihood method to infer *d_N_*/*d_S_* ratio (ω) of *D. melanogaster* protein-coding genes using the genome sequences of five species in *melanogaster* subgroup (*D. melanogaster*, *D. simulans*, *D. sechellia*, *D. yakuba*, and *D. erecta*). The protein-coding sequences were extracted from the alignments of 26 insects, which were obtained from UCSC Genome Browser (http://hgdownload.soe.ucsc.edu/downloads.html). The sequences were further processed by GeneWise (Birney et al., 2004) to remove possible insertions and deletions using the longest isoforms of the corresponding *D. melanogaster* protein sequences as references (FlyBase version r6.15) (Thurmond et al., 2019). The processed sequences were then realigned by PRANK -codon function (Löytynoja, 2014). The conservation scores and coverages of the alignments can be found in Fig. S10. We used codeml in PAML (Yang, 2007) to compute gene-specific ω using M0 model. We removed sequences that have more than 15% of their nucleotides not aligned (gaps) to *D. melanogaster* genes in more than 2 species. To further avoid numeric errors and ensure reasonable estimations, we only retained relatively divergent sequences that are: (1) divergent with dS larger than 0.3, (2) less divergent with dS larger than 0.1 and dN smaller than 0.001 (dS>>dN). At last, there were 12118 genes in total that passed all the criteria and were assigned gene specific ω, containing 6,538,872 amino acids. We also calculated site-specific ω by using likelihood ratio tests (LRT) comparing M7 model against M8 model and M8fix model and M8 model (Yang et al., 2005).

### Rate of adaptive and nonadaptive changes

We recalled all SNPs of 205 inbred lines from the Drosophila Genetic Reference Panel (DGRP), Freeze 2.0 (Huang et al., 2014) (http://dgrp2.gnets.ncsu.edu). We then generated 410 alternative genomes using all monoallelic and bi-allelic SNP data sets. We extracted the coding sequences of *D. melanogaster* genes from the generated alternative genomes, removed all possible insertions and deletions using GeneWise (Birney et al., 2004) as described above. We then align all the coding sequences to their corresponding aligned CDS sequences using PRANK -codon function (Löytynoja, 2014). We removed polymorphisms segregating at frequencies smaller than 5% to reduce possible slightly deleterious mutations (Charlesworth and Eyre-Walker, 2008). In order to avoid possible effects of low divergence between *D. simulans* and *D. melanogaster* (Keightley and Eyre-Walker, 2012), we used *D. yakuba* as outgroup to estimate nonsynonymous polymorphisms (Pn), synonymous polymorphisms (Ps), nonsynonymous substitutions (Dn) and synonymous substitutions (Ds) by MK.pl (Begun et al., 2007; Langley et al., 2012). Similar as Begun et al. (Begun et al., 2007), we only analyzed genes with at least six variants for each of substitutions, polymorphisms, nonsynonymous changes and synonymous changes. We used an extension of MK test, asymptotic MK (Messer and Petrov, 2013; Uricchio et al., 2019), to estimate the proportions of adaptive changes (α). The rate of adaptive changes (ω_a_) was then calculated as ω_a_ = ωα and the rate of non-adaptive changes as ω_na_ = ω - ω_a_. Details of the asymptotic MK test were as following:

1. Classical McDonald–Kreitman test. According to Smith and Eyre-Walker (Smith and Eyre-Walker, 2002), the proportions of adaptive changes for protein-coding genes can be calculated as following:

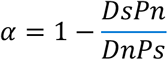 According to this equation, we could estimate the proportion of adaptive changes and carried out classical MK test by applying Fisher’s exact test.
2. Asymptotic estimation of α. A known problem of the classical estimation of α above is the accumulation of slightly deleterious mutations at low frequencies. We therefore used an extension of MK test, asymptotic MK test approach (Messer and Petrov, 2013) to estimate the proportions of adaptive changes. As in original aMK, we defined α(x) as a function of derived allele frequency (x):

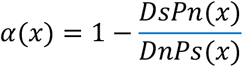

where Pn(x) and Ps(x) are number of non-synonymous and synonymous polymorphisms at frequency x, respectively. However, the original approach may suffer from numeric errors when there were very few polymorphic sites, which is quite common in many of *D. melanogaster* genes. To make the estimations more robust while preserving the same asymptote, we further define Pn (x) and Ps(x) as total number of Pn and Ps above frequency x as described in Uricchio et al (Uricchio et al., 2019). We fitted α(x) to an exponential curve of α(x) ≈ exp(-bx)+c using lmfit (Newville and Stensitzki, 2018) and determined the asymptotic value of α at the limit of x, 1.0. We then estimate the rate of adaptive changes (ω_a_) as

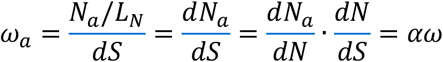

where *N_a_* is the number of adaptive changes and *d_Na_*=*N_a_/L_N_* is the number of adaptive changes per nonsynonymous site. Finally, we calculated the rate of nonadaptive changes (ω_na_) as ω_na_=ω-ω_a_. The final dataset contains 7192 protein-coding genes, with smallest ω_a_ being 0.00 and largest being 1.29.

### Structure-/function- related properties of D. melanogaster proteins

We obtained function-related properties mentioned in the main text as following. We derived *D. melanogaster* gene ages (Kondo et al., 2017; Zhang et al., 2010) for genes that are specific to *Drosophila*, and from GenTree (Shao et al., 2019) for genes that are beyond *Drosophila* clade. We then assigned a pseudo-age to each of the genes. Specifically, there are 11 age groups from “cellular organisms”, assigning to a pseudo age value of 0, to “melanogaster”, assigning a pseudo age value of 10. We downloaded *D. melanogaster* protein-protein interaction (PPI) from STRING database (Szklarczyk et al., 2019). A cut-off of combined score larger than 0.7 was used to retain high confident PPI for further analysis. We then used BSpred (Mukherjee and Zhang, 2011) to predict protein-protein interaction (PPI) sites and DRNApred (Yan and Kurgan, 2017) to predict DNA binding sites. For each protein, we calculated ratios of protein interaction residues (PPI-site ratio) and ratios of DNA binding residues (DNA-site ratio) by dividing total predicted protein interaction sites and DNA binding sites over protein length, respectively. For structure-related properties, we used DeepCNF (Wang et al., 2016) to predict these properties for each gene, including three-state secondary structures (helix, sheet, and coil), structural disorder, relative solvent accessibility (RSA). Further, we calculated the ratios of helix, sheet, helix+sheet, and coil residues of each gene from predicted secondary structures. DeepCNF (Wang et al., 2016) is a deep learning method to capture complex sequence–structure relationships and was proved to have high accuracy in the prediction of protein secondary structures (Yang et al., 2018), structural disorder (Necci et al., 2021) and relative solvent accessibility (Wang et al., 2016). For each gene, we computed intrinsic structural disorder (ISD) and relative solvent accessibility (RSA), as protein-length normalized summations of the probabilities of each residue being disorder and exposed, respectively.

### Gene expression patterns

We downloaded gene expression profile from FlyAtlas2 (Leader et al., 2018). We converted FPKM to TPM by normalizing FPKM against the summation of all FPKMs as following:

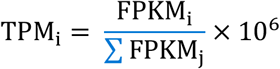

After TPM conversion, we only retained genes with expression level larger than 0.1 TPM for further analysis. We treated male and female whole-body TPM as male and female expression levels. We calculated mean expression level by averaging male and female TPM. We used following Z-score to describe male specificities of *D. melanogaster* genes:

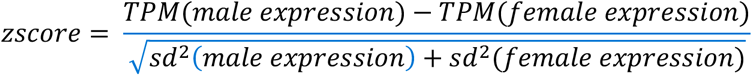

We calculated tissue specificities of genes using tau values (Yanai et al., 2005) based on the expression profiles of 27 different tissues.

### High quality 3D structures of D. melanogaster proteins

We downloaded high-quality structures or structural models of *D. melanogaster* proteins from protein data bank (PDB) (Burley et al., 2019), SWISS-MODEL Repository (Bienert et al., 2017), and MODBASE (Pieper et al., 2011), with descending priorities. For example, if there were 3D structures of the same protein or protein region in multiple databases, we first considered high-resolution structures from PDB; if no structures were found in PDB, we then considered SWISS-MODEL Repository; and at last from MODBASE. In addition, we used blastp (Camacho et al., 2009) to search homologs of each *D. melanogaster* protein against all PDB sequences with an E-value threshold of 0.001. We further carried out comparative structural modeling using RosettaCM (Song et al., 2013) to model high-quality structural models of proteins or protein regions that were not available in PDB, SWISS-MODEL Repository and MODBASE. For each RosettaCM simulation, we used no more than 5 most significant hits from blastp search. For proteins that are in complex forms, we only extracted monomers for further analysis. At last, we obtained 14543 high quality structural models, corresponding to 11284 genes. These structural models contain 2,691,913 unique amino acids, 41.2% of all the residues in genes that were assigned ω.

### Evolutionary rates of different structural/functional sites

We classified amino acids into different classes of structural/functional properties. Specifically, we classified three classes for both ISD and RSA according to the probability of residues being disordered or exposed: ordered or buried (0.00 to 0.33), medium (0.33 to 0.67), disordered or exposed (0.67 to 1.00).

For both PPI and DNA binding, we classified two classes: PPI-site or DNA-site (binding sites), None-PPI or None-DNA (corresponding null sites for PPI or DNA binding). For residues that have 3D structures, we used STRESS (Clarke et al., 2016) to predict putative ligand binding sites and allosteric sites from all the high-quality structures or structural models. We chose STRESS over other programs because it takes both geometry and protein dynamics into account and has been used in genome-wide studies and explained many poorly understood diseases associated variants in humans (Clarke et al., 2016). The allosteric sites were further classified as surface critical or interior critical according to their locations. We then classified these residues into four groups: LIG (ligand binding sites), Surf. Crit. (surface critical sites), Interior Crit. (interior critical sites) and Others (other sites). For each of the site classes, we randomly sampled 100 sequences, each containing 10,000 amino acids. We computed ω, ω_a_, and ω_na_ for the randomly sampled sequences similar to the steps described in the above sections.

### Population genetics of DPGP3 African population

We analyzed 20 high-quality genomes in high latitude South Africa and 30 high-quality genomes in low latitude Ethiopia (Lack et al., 2015). We called all biallelic SNPs and removed SNPs segregating at frequencies smaller than 5% to reduce possible slightly deleterious mutations. Similar to Svetec et al., 2016 (Svetec et al., 2016), for each SNP in both populations, we calculated the fixation index, FST, and used ormidp.test from epitools package in R to perform the odds ratio test for independence. For SNPs at each chromosome arm, we calculated the false discovery rate (FDR) using the Bioconductor *q*-value package (https://github.com/jdstorey/qvalue).

## Acknowledgements

We thank members of the Zhao Lab for helpful discussions, especially Evan Witt for gene expression analysis and Christopher Langer for critically reading an early version of the manuscript.

## Author contribution

J.P. and L.Z. conceived the study. J.P. performed all the analysis with the input from L.Z., N.S. conceptualized the population genetic analysis and helped with the interpretation, J.P. and L.Z. wrote the manuscript.

