## Supplementary information for "Intermolecular interactions drive protein adaptive and co-adaptive evolution at both species and population levels"

**Impact of gene properties on evolution of protein-coding genes in *D. melanogaster***

To uncover the main factors that impact the evolutionary rates of genes, we analyzed 13528 protein-coding genes in *D. melanogaster* using genome data from *melanogaster* subgroup species and *D. melanogaster* population genomics data from 205 inbred lines from *Drosophila* Genetic Reference Panel, Freeze 2.0, DGRP2 (Huang et al., 2014). We applied a maximum likelihood method (Yang, 2007) to compute dN/dS ratio ($\omega$) using the protein-coding sequences of five closely related melanogaster subgroup species (*D. melanogaster*, *D. simulans*, *D. sechellia*, *D. yakuba* and *D. erecta*). We estimated the proportions of adaptive changes ($\alpha$) in each gene by applying an extension of MK test named asymptotic MK (Messer and Petrov, 2013; Uricchio et al., 2019) using *D. simulans* as outgroup. We then calculated the rate of adaptive changes ($\omega_a$) of each gene by multiplying $\omega$ to $\alpha$ ($\omega_a = \alpha\omega$) (Moutinho et al., 2019) using *D. yakuba* as the outgroup species (See methods). The rate of nonadaptive changes can be further calculated by $\omega_{na}=\omega-\omega_a$. Finally, we successfully assigned $\omega$ to 12118 protein-coding genes and $\omega_a$ and $\omega_{na}$ to 7192 genes.

For each of *D. melanogaster* genes subjecting the same analysis pipeline, we further obtained 17 different structural or functional properties (see Methods), which can be further divided into two categories: structure-related properties and function-related properties. Specifically, structure-related properties include the ratio of secondary structures (helix ratio, sheet ratio, helix+sheet ratio, coil ratio), intrinsic structural disorder (ISD), relative solvent accessibility (RSA); while function-related properties include gene pseudo-age, protein length, number of protein-protein interactions (PPI numbers), the ratio of protein-binding sites (PPI-site ratio), the ratio of DNA-binding sites (DNA-site ratio) and gene expression patterns such as male expression level, female expression level, mean expression level, male specificity, and tissue specificity. The properties along with gene-specific protein evolution ($\omega$, $\omega_{a,}$ and $\omega_{na}$) are available in supplementary file S1.

**Molecular interactions contribute to the variations of protein sequence evolution and adaptation**

In order to identify the determinants that drive protein evolution ($\omega$, $\omega_a$ and $\omega_{na}$), we calculated the Pearson's correlations of $\omega$, $\omega_a$ and $\omega_{na}$ with all the structure- and function-related properties. The correlation coefficient (r) and corresponding p-values (p) of each of the properties were listed in Table S1. Interestingly, we observed that for structure-related properties (secondary structure ratios, ISD, and RSA), variation of $\omega$ is dominated by nonadaptive changes ($\omega_{na}$) (Fig. S1). Taking RSA as an example, we observed that RSA strongly correlates with both $\omega$ (r=0.16, p=1e-73) and $\omega_{na}$ (r=0.15, p=3e-35), while weakly correlates with $\omega_a$ (r=0.06, p=1e-6). These correlations suggest that, under the constraints of structure-related properties, relaxation of purifying selection may play a more important role in determining protein evolution. These are in line with previous studies that proteins with less structural constraints, i.e. those harboring more disordered, exposed sites display faster evolutionary and nonadaptive evolutionary rates (Afanasyeva et al., 2018; Moutinho et al., 2019)

However, for function-related properties (gene pseudo-age, protein length, PPI number, PPI-site ratio, DNA-site ratio and gene expression patterns), the importance of $\omega_a$ in shaping protein evolution begins to emerge (Fig. S1). For example, when considering tissue specificity, the correlation efficient (r) of $\omega$ is 0.30 (p=2e-205), while r of $\omega_a$ and $\omega_{na}$ are 0.16 (p=3e-35) and 0.17 (p=2e-42), respectively. In such cases, the correlations of $\omega_a$ and $\omega_{na}$ almost contributed equally to the variation of protein sequence evolutionary rates, $\omega$. Interestingly, among the function-related properties, we found that molecular interactions, i.e., protein interactions, strongly positively correlate with $\omega$, $\omega_a$ and $\omega_{na}$ (Table S1). We also noticed that for molecular interactions, compared to other function-related properties, variations of $\omega_a$ contribute slightly to variations of $\omega$. This could be a result of intercorrelations of molecular interactions and ISD or RSA (Fig. S2), since disordered regions and exposed regions are often responsible for interacting with other molecules (Keskin et al., 2008; Van Der Lee et al., 2014). These results highlight the non-neglected contributions of functional constraints, including molecular interactions, on the adaptive evolution of protein-coding sequences.

**Complex correlations of protein length and male expression level with protein evolutionary rates**

To better clarify and visualize the correlations of $\omega$, $\omega_{a,}$ and $\omega_{na}$ with gene properties in a refined fashion, we divided *D. melanogaster* genes into 15 groups according to the ascending orders of $\omega$ values and compared these properties of different gene groups, while ensuring that each gene group contains the same total number of amino acids (Fig. S3). Overall, for most of the properties being investigated, we observed similar correlations as shown in Table S1. For example, fast evolving genes are relatively young, short, lowly expressed, male or tissue specific, abundant of disordered, exposed residues, excluded in protein-protein network center hubs, and abundant of protein and DNA binding sites.

In contrast to previous observations, we found complex (nonlinear) correlations of ω gene groups with protein length and gene expression levels (Fig. S4). For protein length, our Pearson correlation analysis (Table S1) and a number of previous studies have suggested a strong negative correlation with ω (Lipman et al., 2002; Moutinho et al., 2019). However, we observed that some proteins with the slowest evolutionary rates, i.e. with the smallest ω values, are significantly shorter than other gene groups with intermediate evolutionary rates (Fig. S4A). These include highly conserved genes such as eIF1A (ω=0.0001,148 a.a), rala (ω=0.0001, 201 a.a.), ctp (ω=0.0001, 89 a.a.), and Mlc-c (ω=0.0001, 153 a.a.).

Similar complex correlations were also observed in male expression level and mean expression level (Fig. S4B-C). We found that, when checking male expression level and mean expression level, the gene group that shows the largest mean ω has higher expression than those with intermediate ω. Such U-shape correlations were not observed in female expression levels. Although protein length and mean expression levels of genes are known to be strongly correlated with protein evolutionary rates as listed in Table S1 and also in other references (Drummond et al., 2005; Lipman et al., 2002; Zhang and Yang, 2015), fast evolving genes can also be moderately or highly expressed, especially in male *D. melanogaster*. For example, many seminal fluid proteins show high ω values and are highly expressed, such as Sfp60F (ω=0.77, 82 a.a.), EbpII (ω=0.68, 66 a.a), Acp36DE (ω=0.68, 912 a.a.,), and Dup99B (ω=0.63, 54 a.a.). These proteins evolve at very fast rates (Begun and Lindfors, 2005; Swanson et al., 2001), contain a various range of amino acids (54 in Dup99B to 912 in Acp36DE), and are moderately or highly expressed in male *D. melanogaster* (TPM ranging from 440 for Acp36DE to 3189 for Sfp60F), lowly expressed in female (TPM all around 1, presumably in spermatheca). We listed all the genes and protein length and expression levels in each ω gene group, which can be found in supplementary file S2.

Since tissue specificity and male specificity both strongly correlate with ω, $\omega_a$, and $\omega_{na}$ (Table S1), we asked whether male specificity would be a redundant property compared to tissue specificity to indicate protein evolution due to the complex correlations of male expression levels. To answer this question, we classified *D. melanogaster* genes into 15 groups according to ascending values of male specificity. We then did a similar classification to classify all the genes into 15 groups according to ascending values of tissue specificity (Fig. S5). As expected, we found that tissue specificity positively correlates with ω, $\omega_a$ and $\omega_{na}$. However, we observed complex correlations for male specificity gene groups. Specifically, the gene group with the lowest male specificity shows significantly higher ω, $\omega_a$ and $\omega_{na}$ than its following gene group (Fig. S5). This could be a result of fast evolving female-biased genes (Yang et al., 2016) included in this gene group.

Although our results are in agreement with previous studies on the factors driving protein sequence evolution (Zhang and Yang, 2015), we showed some complex correlations between ω, ωa and ωna and protein length and male specificity (Fig. S8-S9, supplement file S2). These complex correlations suggest caveat exists when we looked at protein length and gene expression levels. For example, gene expression level was proved to be a major determinant (Zhang and Yang, 2015) through mechanisms such as the pressure for translational robustness, i.e., robustness to translational missense errors (Drummond et al., 2005). Previous studies have revealed that male biased or female biased genes can be fast evolving (Yang et al., 2016). While on the other hand, many male biased genes can be highly expressed in testis, which results in a complex correlation between protein sequence evolutionary rate and male expression level or even mean expression level of D. melanogaster. The unique evolutionary property of these male-biased or specific genes could be caused by the unique transcriptional scanning mechanism in testis (Xia et al., 2020). We propose that tissue specificity might be a better quantity when considering the impact of gene expression profile on protein sequence evolution in *D. melanogaster*. In addition to male expression level, a similar complex correlation was observed for protein length. It has been the notion that short proteins tend to evolve faster than long proteins, which may be biologically relevant or a byproduct of other factors such as selection on buried and exposed sites (Moutinho et al., 2019). Here, we demonstrated that, in D. melanogaster, although protein length is strongly negatively correlated with protein sequence evolutionary rate, genes that have the slowest evolutionary rates tend to be relatively short. This could be caused by the fact that under essential functional constraints, genes can undergo strong purifying selections, while essential genes such as secreted proteins are constrained to be smaller, and that essential genes could be shorter than other genes (Chen et al., 2020).

**Table S1.** Pearson correlation coefficients between $\omega$, $\omega_a$ and $\omega_{na}$ and gene properties [a]

| Categories | Properties | $\omega$ | | $\omega_a$ | | $\omega_{na}$ | |
|---|---|---|---|---|---|---|---|
| Function-related properties | Gene age | 0.55 | (0) | 0.34 | (7e-154) | 0.41 | (2e-224) |
| | Protein length [b] | -0.22 | (2e-136) | -0.13 | (1e-25) | -0.30 | (2e-142) |
| | Mean expression [b] | -0.21 | (2e-109) | -0.11 | (2e-18) | -0.11 | (1e-18) |
| | Male expression [b] | -0.06 | (5e-10) | 0.00 | (8e-1) | -0.03 | (6e-2) |
| | Female expression [b] | -0.29 | (2e-205) | -0.17 | (2e-42) | -0.16 | (3e-35) |
| | Male specificity | 0.21 | (2e-104) | 0.12 | (3e-22) | 0.10 | (3e-14) |
| | Tissue Specificity | 0.30 | (4e-226) | 0.18 | (1e-45) | 0.19 | (3e-48) |
| | PPI number [b] | -0.28 | (1e-217) | -0.14 | (4e-29) | -0.19 | (4e-58) |
| | PPI-site ratio | 0.14 | (1e-50) | 0.05 | (7e-6) | 0.10 | (2e-16) |
| | DNA-site ratio | 0.25 | (8e-164) | 0.12 | (3e-23) | 0.23 | (3e-79) |
| Structure-related properties | Helix ratio | -0.05 | (4e-7) | -0.01 | (3e-1) | -0.05 | (4e-5) |
| | Sheet ratio | -0.04 | (2e-5) | 0.00 | (9e-1) | -0.01 | (3e-1) |
| | Helix+sheet ratio | -0.09 | (3e-25) | -0.02 | (1e-1) | -0.07 | (1e-9) |
| | Coil ratio | 0.10 | (8e-27) | 0.01 | (2e-1) | 0.08 | (8e-11) |
| | ISD | 0.17 | (8e-82) | 0.04 | (1e-3) | 0.12 | (2e-24) |
| | RSA | 0.16 | (7e-87) | 0.06 | (1e-6) | 0.15 | (3e-35) |
| Protein evolution | $\omega$ | 1.00 | (0) | 0.65 | (0) | 0.78 | (0) |
| | $\omega_a$ | 0.65 | (0) | 1.00 | (0) | 0.03 | (7e-3) |
| | $\omega_{na}$ | 0.78 | (0) | 0.03 | (7e-3) | 1.00 | (0) |

[a] Pearson correlation coefficient R were listed along with corresponding P-values in parentheses.
[b] To better estimate the correlations for sequence length, expression levels and PPI numbers, we used logarithmic scales rather than absolute values, which could vary dramatically from near zero to thousands.
Abbreviations in this table: ISD, intrinsic structural disorder; RSA, relative solvent accessibility; PPI number, protein-protein interaction number; PPI-site ratio, ratio of protein-protein interaction sites; DNA-site ratio, ratio of DNA-binding sites.

**Table S2**. Detailed descriptions of 19 protein properties

| Categories | Properties | Descriptions |
|---|---|---|
| Function-related properties | Gene age | Pseudogene ages, with index 10 being D. melanogaster specific and 0 being existed in all living cells, see *Structure-/function- related properties of D. melanogaster proteins* in Material and Methods |
| | Protein length | Number of amino acids in the longest isoform |
| | Mean expression | Mean of the TPM levels in female whole body and male whole body |
| | Male expression | TPM levels in male whole body |
| | Female expression | TPM levels in female whole body |
| | Male specificity | zscore in section *Gene expression patterns* in *Methods* |
| | Tissue Specificity | tau in section *Gene expression patterns* in Material and Methods |
| | PPI number [b] | Number rof protein patterners from STRING database |
| | PPI-site ratio | Fraction of residues inside the gene that are involved in protein-protein interactions |
| | DNA-site ratio | Fraction of residues inside the gene that are involved in protein-protein interactions |
| Structure-related properties | Helix ratio | Fraction of helix residues inside the gene |
| | Sheet ratio | Fraction of beta residues inside the gene |
| | Helix+sheet ratio | Fraction of helix and beta residues inside the gene |
| | Coil ratio | Fraction of random coil residues inside the gene |
| | ISD | Intrinsic structural disorder, probability of a gene being intrinsically disordered |
| | RSA | Relative solvent accessibility, probability of a gene being solvent accessible |
| Protein evolution | $\omega$ | Evolutionary rate |
| | $\omega_a$ | Adaptation rate |
| | $\omega_{na}$ | Nonadaptation rate |

**Table S3**. Protein families in fast-adaptive proteins

| Term | Count | Benjamini P-value |
| --- | --- | --- |
| Peptidase S1 | 45 | 9.7E-12 |
| Trypsin-like cysteine/serine peptidase domain | 45 | 1.1E-11 |
| Peptidase S1A, chymotrypsin-type | 36 | 4.4E-8 |
| Peptidase S1, trypsin family, active site | 31 | 1.8E-7 |
| 7TM chemoreceptor | 14 | 4.1E-5 |
| Protein of unknown function DUF1091 | 17 | 8.2E-4 |
| Chitin binding domain | 17 | 1.7E-3 |

**Table S4**. Go analysis of fast-adaptive genes

| **Go Term** | **#** | **Fold Enrichment** | **Benjamini P-value** |
|---|---|---|---|
| serine-type endopeptidase activity | 45 | 3.9 | 1.9E-12 |
| multicellular organism reproduction | 43 | 4.2 | 2.9E-12 |
| proteolysis | 58 | 3.0 | 1.7E-11 |
| sensory perception of bitter taste | 14 | 6.8 | 1.1E-5 |
| response to carbon dioxide | 14 | 7.0 | 1.1E-5 |
| chemosensory behavior | 14 | 6.3 | 2.3E-5 |
| sweet taste receptor activity | 14 | 5.9 | 6.5E-5 |
| taste receptor activity | 14 | 5.5 | 9.8E-5 |
| olfactory receptor activity | 19 | 3.9 | 1.1E-4 |
| chitin metabolic process | 17 | 4.3 | 1.9E-4 |
| sensory perception of taste | 13 | 5.0 | 8.0E-4 |
| male courtship behavior | 16 | 3.7 | 1.8E-3 |
| chitin binding | 17 | 3.4 | 2.3E-3 |
| negative regulation of transposition | 6 | 13.6 | 2.8E-3 |
| procollagen-proline 4-dioxygenase activity | 7 | 6.6 | 2.5E-2 |
| sensory perception of pain | 42 | 1.7 | 3.7E-2 |
| oxidoreductase activity, acting on single donors with incorporation of molecular oxygen, incorporation of two atoms of oxygen | 7 | 5.9 | 4.0E-2 |
| dopamine:sodium symporter activity | 4 | 16.2 | 4.5E-2 |

**Table S5.** Sub-clusters in the PPI network of fast-adaptive proteins

| # sub-cluster | Genes |
|---|---|
| Sub-cluster 1 | CG12729, p53, PH4alphaNE2, Tango1, CG34130, Fadd, CG30054, spz, CG17097, Spn28F, IKKbeta, Acp53Ea, CG14074, wek, CG31642, CG32413, CG10909, Hrd3, dgrn, Qsox3, Rel, CG12477, PGRP-LC, Ebp, CG9997, CG31496, CG6168, CG14443, CG32373, Acp53C14b, Obp56i, Spn77Bb, Rad60, CG18258, CG5381, Sfp26Ad, Herp, Dronc, CG34129, CG15445, APC7, Send2, Acp62F, CG14322, l(2)03659, Acp53C14a, Toll-4, CG15784, CG7386, CG7824, CG5111, Qsox4, Daxx, CG7362, p38c |
| Sub-cluster 2 | Mst33A, CG2267, CG15631, CG5217, CG31948, salto, CG14835, vrs, CG10177, spaw, CG4712, CG17666, h-cup, c-cup, CG5614, CG6304, CG32081 |
| Sub-cluster 3 | spn-E, Tdrd3, armi, nos, squ, swa, osk, rhi, Fancm, r2d2, Snm1, Dcr-2, tej, tapas, aub |
| Sub-cluster 4 | Nuf2, Klp3A, Mzt1, Grip84, Mps1, MCPH1, Grip128, cmet, Cap-H2, Klp59D, cnn, Nnf1b, Grip75, Cap-G |
| Sub-cluster 5 | ItgaPS4, Btnd, CG31358, CG32750, CG14736, vlc |
| Sub-cluster 6 | Cul6, CG5003, CG43089, CG31633, CG34025 |
| Sub-cluster 7 | CG43896, CG6296, CG5883, CG17826, CG42397 |

**Table S6**. Go term enrichment of proteins in sub-cluster 1.

| Go Term | Count | Fold Enrichment | Benjamini P-value | Genes |
|---|---|---|---|---|
| multicellular organism reproduction | 18 | 16.0 | 1.9E-14 | Sfp26Ad, Ebp, Obp56i, Spn28F, Spn77Bb, PH4alphaNE2, Acp53C14a, Acp53C14b, Acp53Ea, Acp62F, CG17097, CG31413, CG34129, CG34130, CG6168, CG6690, CG9997 |
| positive regulation of antibacterial peptide biosynthetic process | 4 | 76.4 | 1.4E-3 | Fadd, IKKbeta, PGRP-LC, Rel |
| peptidoglycan recognition protein signaling pathway | 4 | 50.9 | 3.3E-3 | Fadd, IKKbeta, PGRP-LC, Rel |
| Toll signaling pathway | 4 | 26.2 | 1.9E-2 | IKKbeta, Rel, spz, wek |
| positive regulation of defense response to virus by host | 3 | 68.7 | 2.7E-2 | Fadd, IKKbeta, Rel |
| defense response to Gram-negative bacterium | 5 | 10.6 | 3.3E-2 | Fadd, IKKbeta, Rel, PGRP-LC, CG6168 |
| immune response | 4 | 17.0 | 3.8E-2 | Fadd, PGRP-LC, Rel, spz |
| activation of cysteine-type endopeptidase activity involved in apoptotic process | 3 | 43.0 | 4.4E-2 | p53, Dronc, Fadd |

**Table S7.** Go term enrichment of proteins in sub-cluster 3.

| Go Term | Count | Fold Enrichment | Benjamini P-value | Genes |
|---|---|---|---|---|
| negative regulation of transposition | 4 | 266.6 | 2.8E-5 | spn-E, aub, tej, CG8920 |
| regulation of pole plasm oskar mRNA localization | 4 | 122.2 | 1.7E-4 | spn-E, aub, armi, swa |
| RNA interference | 4 | 91.6 | 2.7E-4 | spn-E, aub, armi, Dcr-2 |
| targeting of mRNA for destruction involved in RNA interference | 3 | 439.8 | 3.9E-4 | spn-E, aub, Dcr-2 |
| oogenesis | 6 | 14.0 | 6.2E-4 | spn-E, aub, armi, osk, nos, squ |
| dorsal appendage formation | 4 | 54.3 | 6.7E-4 | spn-E, aub, armi, squ |
| segmentation | 3 | 20.0 | 1.5E-3 | aub, osk, nos |
| pole plasm protein localizaiton | 3 | 20.0 | 1.5E-3 | aub, armi, osk |
| karyosome formation | 3 | 20.0 | 6.2E-3 | osk, nos, swa |
| anterior/posterior axis specification, embryo | 3 | 20.0 | 1.8E-2 | osk, nos, swa |
| piRNA biosynthetic process | 2 | 13.3 | 3.9E-2 | tej, CG8920 |
| oocyte maturation | 2 | 13.3 | 4.7E-2 | spn-E, aub |
| siRNA loading onto RISC involved in RNA interface | 2 | 13.3 | 4.7E-2 | Dcr-2, r2d2 |

**Table S8**. Go term enrichment of proteins in sub-cluster 4.

| Go Term | Count | Fold Enrichment | Benjamini P-value | Genes |
|---|---|---|---|---|
| interphase microtubule nucleation by interphase microtubule organizing center | 4 | 349.1 | 7.1E-6 | Grip75, Grip84, Grip128, CG42787 |
| mitotic nuclear division | 6 | 33.9 | 1.2E-5 | Grip75, Grip84, Grip128, Klp3A, MCPH1, cnn |
| centrosome organization | 4 | 56.1 | 7.6E-4 | Grip75, Grip84, Grip128, cnn |
| Homestasis (R-DME-983189) | 3 | 184.5 | 4.6E-5 | Klp3A, Klp59D, cmet |
| mitotic metaphase plate congression | 3 | 181.3 | 1.6E-3 | Grip75, Grip84, Grip128 |
| microtubule nucleation | 3 | 138.6 | 2.3E-3 | Grip75, Grip84, Grip128 |
| mitotic spindle assembly | 3 | 81.3 | 5.6E-3 | Grip75, Grip84, CG42787 |
| chromosome segregation | 3 | 65.5 | 7.5E-3 | Klp3A, Nuf2, Nnf1b |
| female meiosis chromosome segregation | 3 | 39.3 | 1.8E-2 | Klp3A, Mps1, cnn |
| microtubule-based movement | 3 | 28.1 | 3.1E-2 | Grip75, Grip84, CG42787 |
| spindle assembly involved in female meiosis II | 2 | 392.7 | 3.1E-2 | Grip75, Grip128 |
| centrosome duplication | 3 | 21.8 | 4.1E-2 | Grip75, Grip84, Grip128 |

**Table S9**. Go term enrichment of proteins in sub-cluster 5.

| Go Term | Count | Fold Enrichment | Benjamini P-value | Genes |
|---|---|---|---|---|
| nitrogen compound metabolic process | 2 | 343.6 | 4.7E-2 | Btnd, CG32750 |

**Table S10.** Go term enrichment of proteins in sub-cluster 6.

| Go Term | Count | Fold Enrichment | Benjamini P-value | Genes |
|---|---|---|---|---|
| SCF-dependent proteasomal ubiquitin-dependent protein catabolic process | 3 | 140.4 | 9.6E-4 | CG31633, CG34025, CG5003 |
| protein ubiquitination | 3 | 45.2 | 4.6E-3 | CG31633, CG34025, CG5003 |
| ubiquitin-dependent protein catabolic process | 2 | 61.9 | 7.5E-2 | CG11261, CG43089 |

**Table S11.** Go term enrichment of proteins in sub-cluster 7.

| Go Term | Count | Fold Enrichment | Benjamini P-value | Genes |
|---|---|---|---|---|
| chitin metabolic process | 4 | 88.9 | 8.4E-6 | CG17826, CG42397, CG43896, CG5883 |

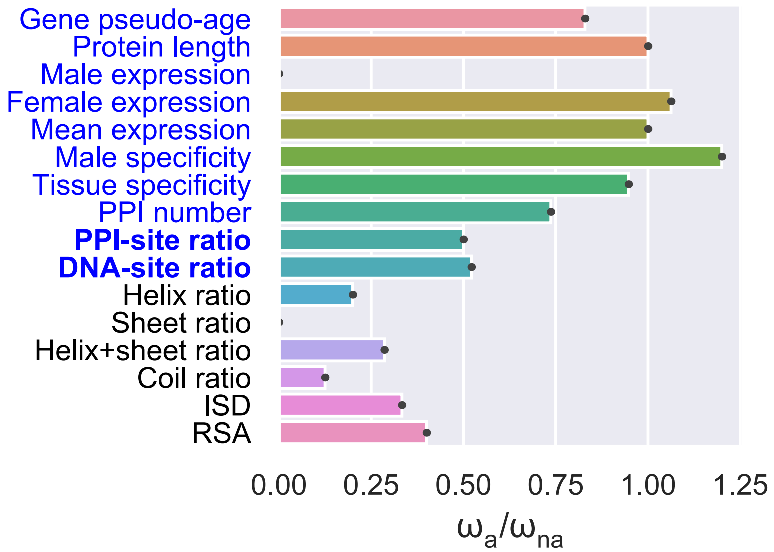


**Fig. S1**. Contributions of adaptation rates ($\omega_a$) and non-adaptation rates ($\omega_{na}$) to the variation of evolutionary rates ($\omega$) of protein-coding sequences in *D. melanogaster*. Under the constraints of structure-related properties (black tick label), variation of $\omega$ is dominated by $\omega_{na}$. While for function related properties (blue tick label), $\omega_a$ contributes almost equally as $\omega_{na}$ to variation of $\omega$. In addition, for molecular interactions, i.e., PPI-site ratio and DNA-site ratio, the role of $\omega_a$ in determine the variation of $\omega$ is non-neglected.

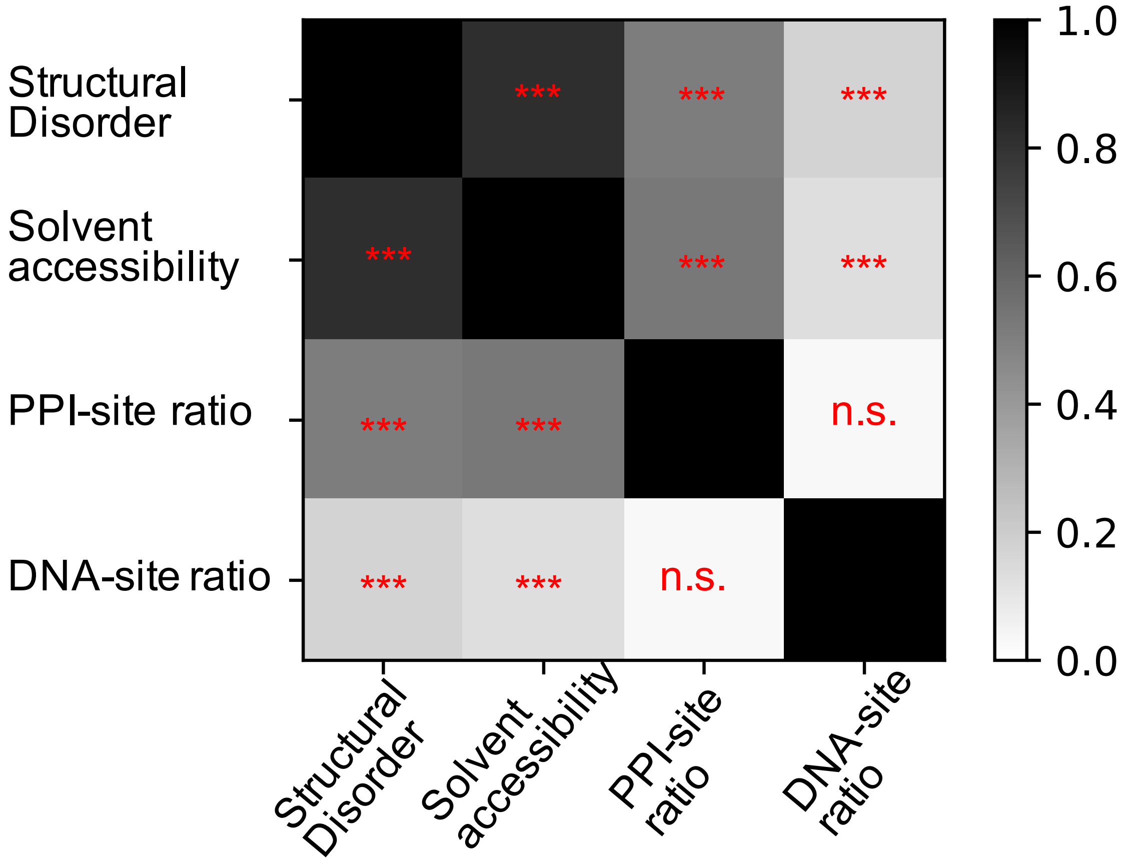


**Fig. S2.** Pearson correlations between structural disorder, solvent accessibility and molecular interactions. Significance level *** (P<1e-80).

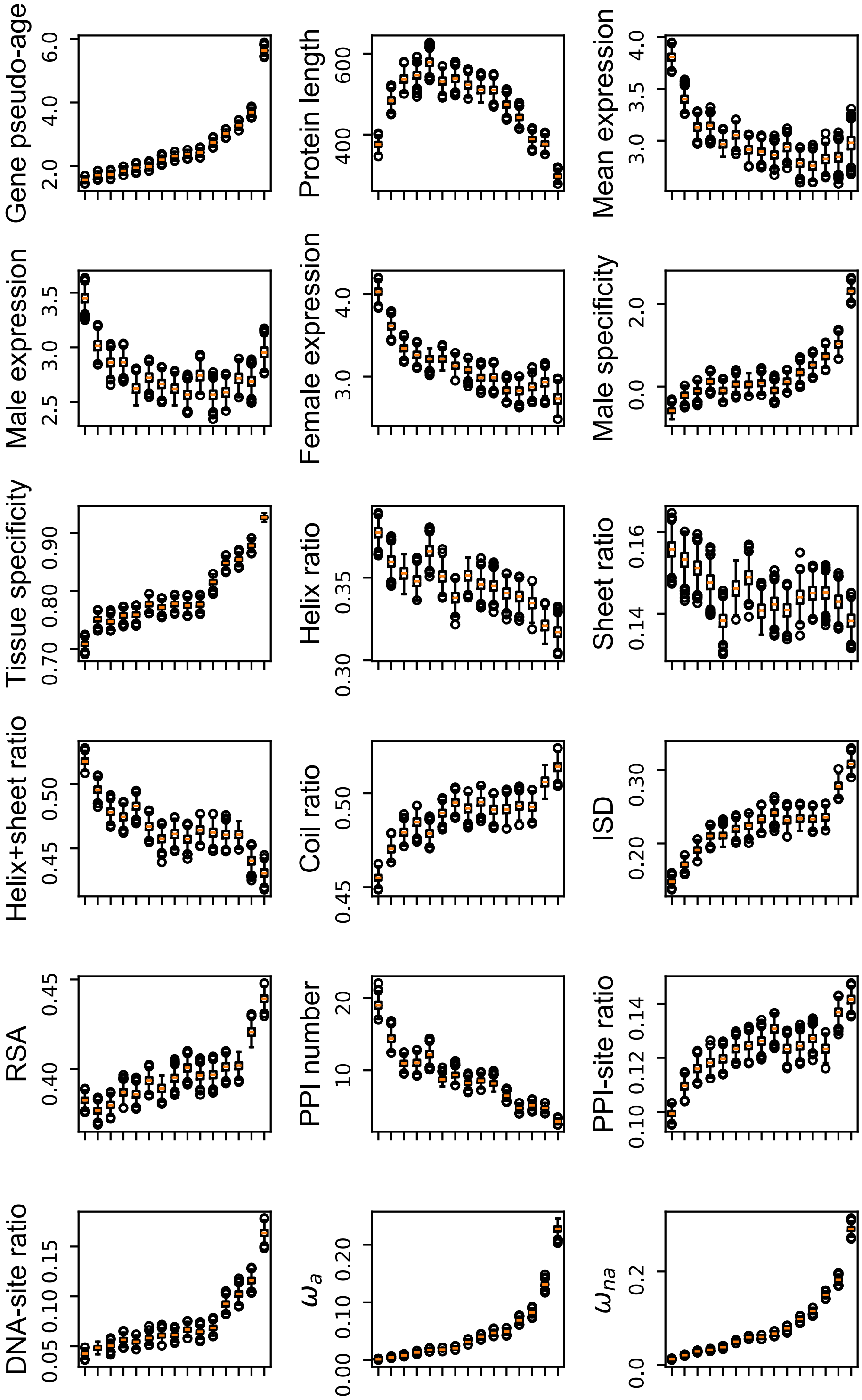


**Fig. S3**. Gene properties in each gene groups classified by the ascending order of evolutionary rates.

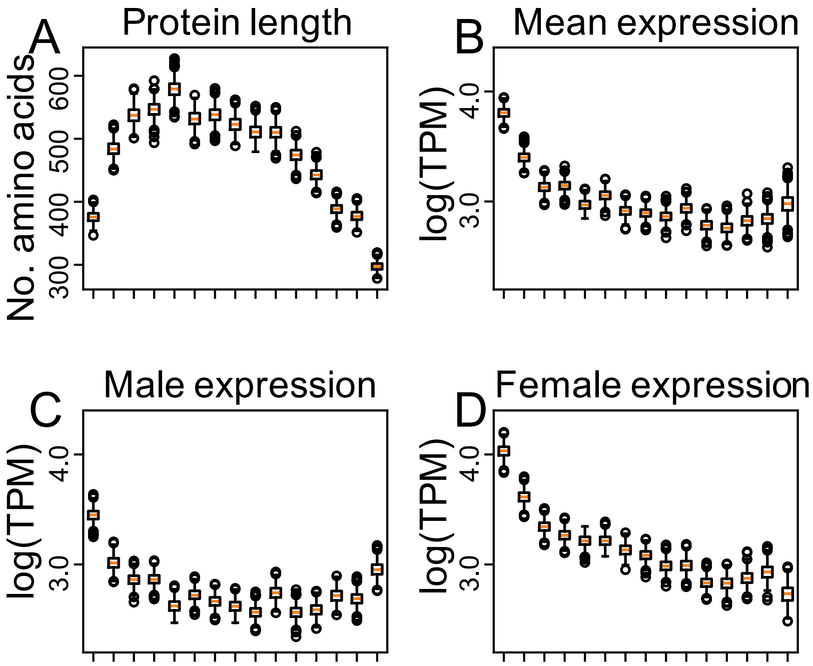


**Fig. S4**. Protein length (A), mean (B), male (C) and female expression (D) levels in each gene groups divided by ascending *ω* values. Values for each gene group and each gene property were computed through 1,000 bootstrapping steps. Obvious complex U-shaped correlations with *ω* were observed for protein length (A) and male expression level (C).

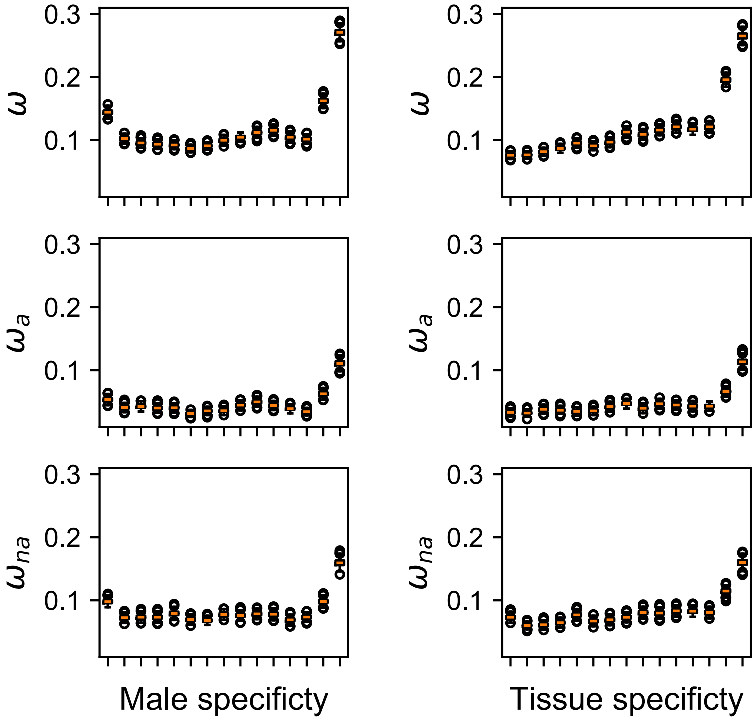


**Fig. S5**. Protein evolutionary rates (ω, $\omega_a$ and $\omega_{na}$) in genes with different male specificity (left panel) and tissue specificity (right panel). Complex correlations were observed for male specificity, i.e., gene groups with the lowest male specificity shows higher ω, $\omega_a$ and $\omega_{na}$ than the second lowest gene group.

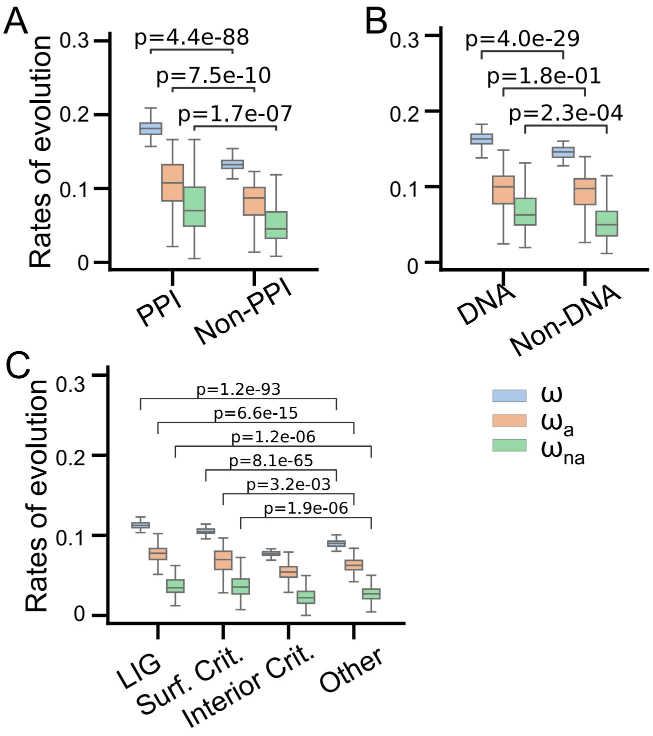


**Fig. S6**. Evolutionary rates of putative binding sites, including protein-protein interaction sites (A), DNA binding sites (B), and ligand binding sites (C), are higher than corresponding none sites.

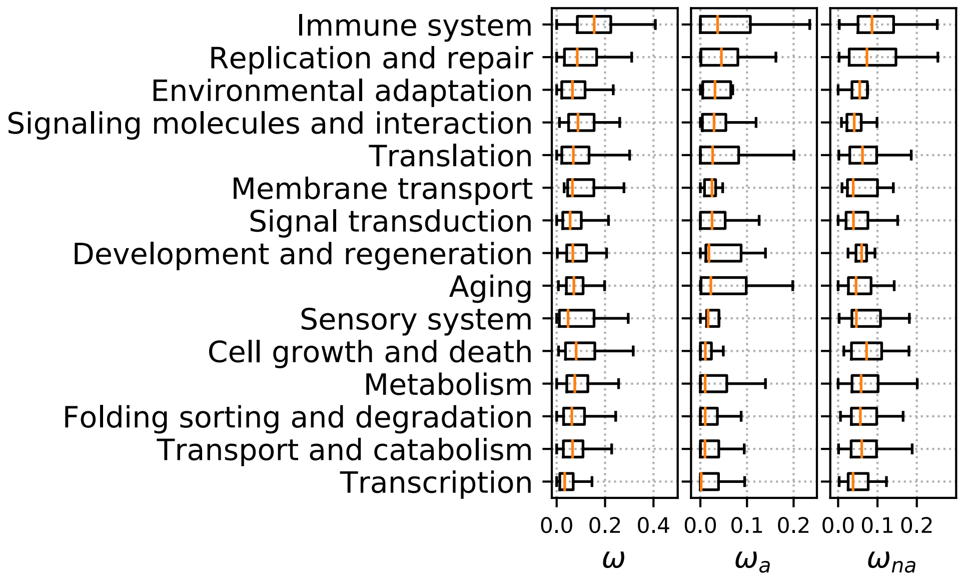


**Fig. S7.** Functional pathways that are associated with immunity and environmental adaptation have higher adaptive evolutionary rates than other pathways.

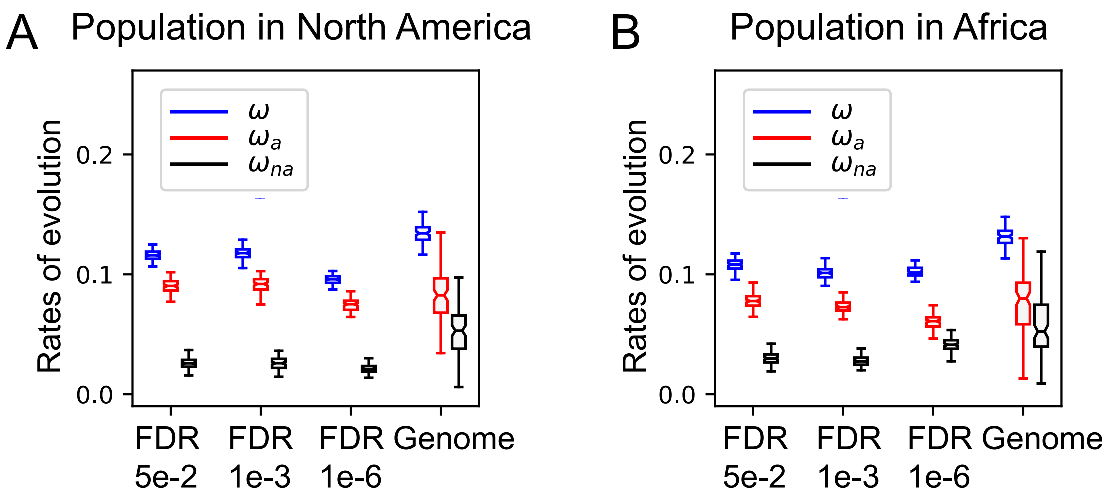


**Fig. S8.** Evolutionary rates of significantly differentiated SNPs in (A) North American population and (B) African population.

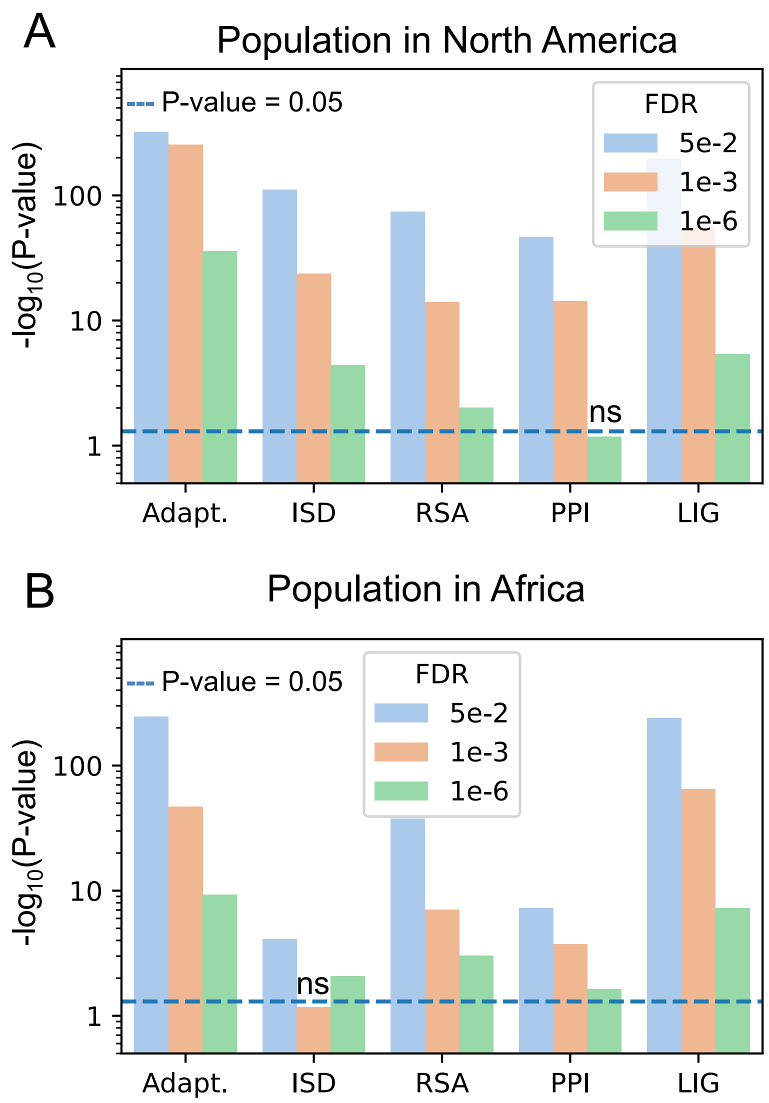


**Fig. S9.** Fisher exact test of enrichment of significantly differentiated nonsynonymous SNPs for (A) North American population and (B) African population. Differentiated nonsynonymous SNPs of both the two populations are enriched in long term adaptive evolution (adaptive evolution, Adapt.), intrinsic disorder regions (ISD), solvent accessible regions (RSA), protein-protein interactions (PPI), and ligand-binding interactions (LIG). We marked two situations where p-values were smaller than 0.05 with "ns": enrichment of North American differentiated nonsynonymous SNPs (FDR 1e-6) in protein-protein interaction (PPI) sites (p-value = 0.07) and enrichment of African differentiated nonsynonymous SNPs (FDR 1e-3) in intrinsic disordered (ISD) regions (p-value = 0.07). In other cases, p-values were all smaller than 0.05.

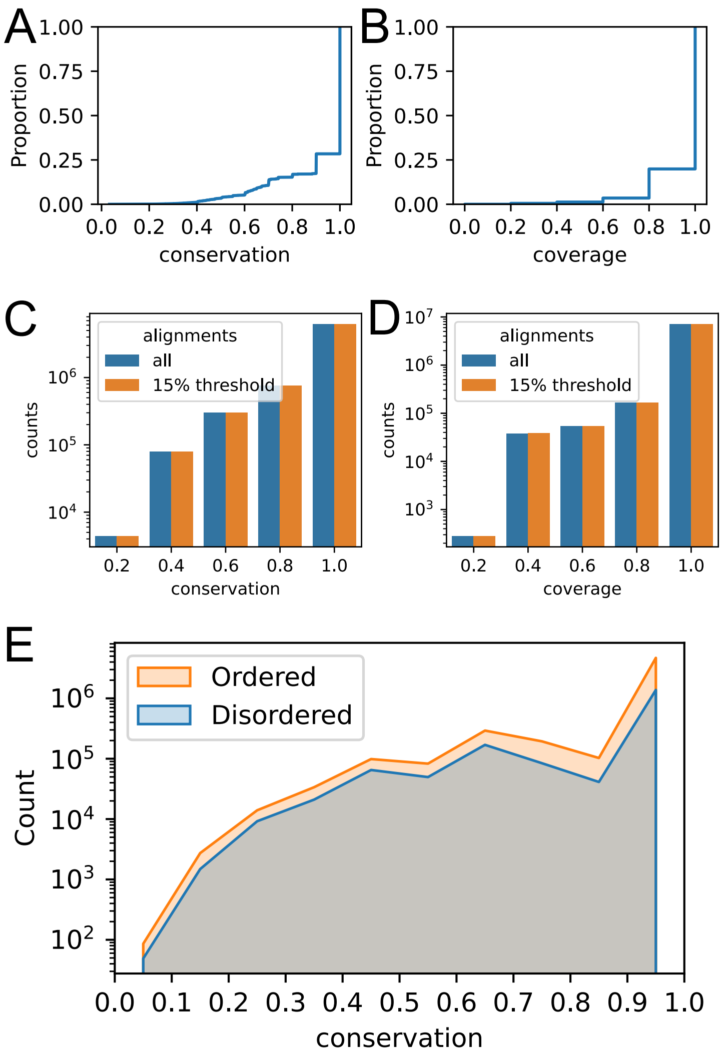


**Fig. S10**. Conservation scores and coverages of protein alignments. The accumulative distribution of conservation scores (A) and coverages (B) of protein alignments used in our study. The counts of sites of all alignments derived from UCSC Genome browser (blue bar, C and D) and after removing alignments that have more than 15% of the sites removed (orange bar, C and D). (E) Disordered regions (blue line) share similar distributions of conservations as ordered regions (orange line).
